# An atlas of cell types in the mammalian epididymis and vas deferens

**DOI:** 10.1101/2020.01.24.918979

**Authors:** Vera D. Rinaldi, Elisa Donnard, Kyle J. Gellatly, Morten Rasmussen, Alper Kucukural, Onur Yukselen, Manuel Garber, Upasna Sharma, Oliver J. Rando

**Affiliations:** Department of Biochemistry and Molecular Pharmacology, University of Massachusetts Medical School, Worcester, MA 01605, USA; Department of Bioinformatics and Integrative Biology, University of Massachusetts Medical School, Worcester, MA 01605, USA; Department of Virus and Microbiological Special Diagnostics, Statens Serum Institut, Copenhagen, Denmark; Program in Molecular Medicine, University of Massachusetts Medical School, Worcester, MA 01605, USA; Department of Molecular, Cell and Developmental Biology, University of California Santa Cruz, Santa Cruz, CA 95064, USA

## Abstract

Following spermatogenesis in the testis, mammalian sperm continue to mature over the course of approximately 10 days as they transit a long epithelial tube known as the epididymis. The epididymis is comprised of multiple segments/compartments that, in addition to concentrating sperm and preventing their premature activation, play key roles in remodeling the protein, lipid, and RNA composition of maturing sperm. In order to understand the complex roles for the epididymis in reproductive biology, we generated a single cell atlas of gene expression from the murine epididymis and vas deferens. We recovered all the key cell types of the epididymal epithelium, including principal cells, clear cells, and basal cells, along with associated support cells that include fibroblasts, smooth muscle, macrophages and other immune cells. Moreover, our data illuminate extensive regional specialization of principal cell populations across the length of the epididymis, with a substantial fraction of segment-specific genes localized in genomic clusters of functionally-related genes. In addition to the extensive region-specific specialization of principal cells, we find evidence for functionally-specialized subpopulations of stromal cells, and, most notably, two distinct populations of clear cells. Analysis of ligand/receptor expression reveals a network of potential cellular signaling connections, with several predicted interactions between cell types that may play roles in immune cell recruitment and other aspects of epididymal function. Our dataset extends on existing knowledge of epididymal biology, and provides a wealth of information on potential regulatory and signaling factors that bear future investigation.

## INTRODUCTION

In mammals, the production of sperm that are competent to fertilize the egg requires multiple organs that together constitute the male reproductive tract. A common characteristic among these is the presence of highly specialized somatic cells that support the development and maturation of gametes. For instance, testicular spermatogenesis requires multiple functionally-distinct cells that support the development and function of sperm. These include several cell types that are intimately associated with the developing sperm in the testis, most notably the Sertoli cells, which envelop and provide support to developing sperm, and the Leydig cells, which are located in the testicular interstitial space and are responsible for testosterone biosynthesis.

Following the process of testicular spermatogenesis and spermiogenesis, sperm are morphologically mature, but do not yet exhibit forward motility and are incapable of fertilization. Sperm maturation continues after sperm exit the testis, at which point they enter a long convoluted tube known as the epididymis where they continue to mature over the course of ∼10 days in the mouse. Epididymal transit has long been understood to be essential for male fertility (Bedford, 1967; Orgebin-Crist, 1967; Young, 1931), and sperm undergo extensive molecular and physiological changes during this process (Cooper, 2015; Gervasi and Visconti, 2017). Sperm are concentrated as they proceed from caput (proximal) to corpus and then to the cauda (distal) epididymis, where they can be stored for prolonged periods in an inactive state thanks to the ionic microenvironment of the cauda epididymal lumen. In addition, sperm are extensively remodeled during epididymal transit. Most notably, sperm proteins –including, but not limited to, protamines – become extensively disulfide-crosslinked during epididymal maturation, and this is thought to mechanically stabilize various sperm structures (Balhorn, 1982; Calvin and Bedford, 1971). At the sperm surface, lipid remodeling events include cholesterol removal and an increase in levels of polyunsaturated fatty acids with an accompanying increase in membrane fluidity (Saez et al., 2011), while maturation of the glycocalyx involves extensive remodeling of accessible sugar moieties (Tecle and Gagneux, 2015; Tulsiani, 2006). In addition, a heterogeneous population of extracellular vesicles collectively known as epididymosomes deliver proteins and small RNAs – some of which play functional roles in fertilization and early development – to maturing sperm (Caballero et al., 2013; Conine et al., 2018; Krapf et al., 2012; Martin-DeLeon, 2015; Reilly et al., 2016; Sharma et al., 2016; Sharma et al., 2018; Sullivan et al., 2007; Sullivan and Saez, 2013).

The organ responsible for this ever-expanding diversity of functions is a long tube on the order of 1-10 meters long when extended, depending on the species – of pseudo-stratified epithelium. The functions of the epididymis differ extensively along its length (Domeniconi et al., 2016; Turner et al., 2003), as for example many genes such as *Rnase10* and *Lcn8* are expressed almost exclusively in the proximal epididymis, most notably including the testis-adjacent section of the caput epididymis known as the “initial segment” (Hsia and Cornwall, 2004; Jelinsky et al., 2007; Jervis and Robaire, 2001; Johnston et al., 2005; Turner et al., 2003). Connective tissue septa separate the mouse epididymis into ten anatomically-defined segments (this number varies in other mammals) (Turner et al., 2003), with early microarray studies revealing a wide variety of gene expression profiles across these ten segments, and defining roughly six distinct gene expression domains across the epididymis (Johnston et al., 2005). Dramatic gradients of gene expression along the epididymis are also observed in many other mammals, including rat (Jelinsky et al., 2007; Jervis and Robaire, 2001), boar (Guyonnet et al., 2009), and human (Thimon et al., 2007). In addition to the variation in gene regulation along the length of the epididymis, the epididymal epithelium at any point along the path is comprised of multiple morphologically- and functionally-distinct cell types, including principal cells, basal cells, and clear cells (Breton et al., 2016).

Here, we sought to further explore the cellular makeup and gene expression patterns across this relatively understudied organ. Over the past few years, microfluidic-based cell isolation and molecular barcoding strategies, coupled with continually-improving genome-wide deep sequencing methods, have facilitated the rise of ultra-high throughput analysis of gene expression in thousands of individual cells from a wide range of tissues (Cao et al., 2017; Farrell et al., 2018; Han et al., 2018; Ramskold et al., 2012). Using droplet-based (10X Chromium) single cell RNA-Seq, we profiled gene expression in 8,880 individual cells from across the epididymis and the vas deferens. Our data confirm and extend prior studies of segment-restricted gene expression programs, and reveal that these region-specific programs are largely driven by principal cells. We find several novel cell subpopulations, most notably including an Amylin-positive clear cell subtype, and predict intercellular signaling networks between the various cell types. Together, these data provide an atlas of cell composition across one of the least-studied organs in the body, and provide a wealth of molecular hypotheses for future efforts in reproductive physiology.

## RESULTS

### Dataset generation and overview

We set out to characterize the regulatory program across the post-testicular male reproductive tract, focusing on four tissue samples – the caput, corpus, and cauda epididymis, as well as the vas deferens (**Figure 1A**). We first generated a baseline dataset via traditional RNA-Seq profiling for at least 10 individual samples of the caput, corpus, cauda and vas deferens (**Table S1**). Our bulk RNA-Seq dataset confirms the expected region-specific gene expression previously observed in many species (**Figure 1B, Table S1**), and recapitulates, at coarser anatomical resolution but improved genomic resolution, prior microarray analyses of the murine epididymis (Johnston et al., 2005) (**Figure 1-figure supplement 1**). Furthermore, we include below a more detailed analysis of gene expression in the vas deferens, a tissue which is not often included in published studies of the epididymis transcriptome.

**Figure 1.**
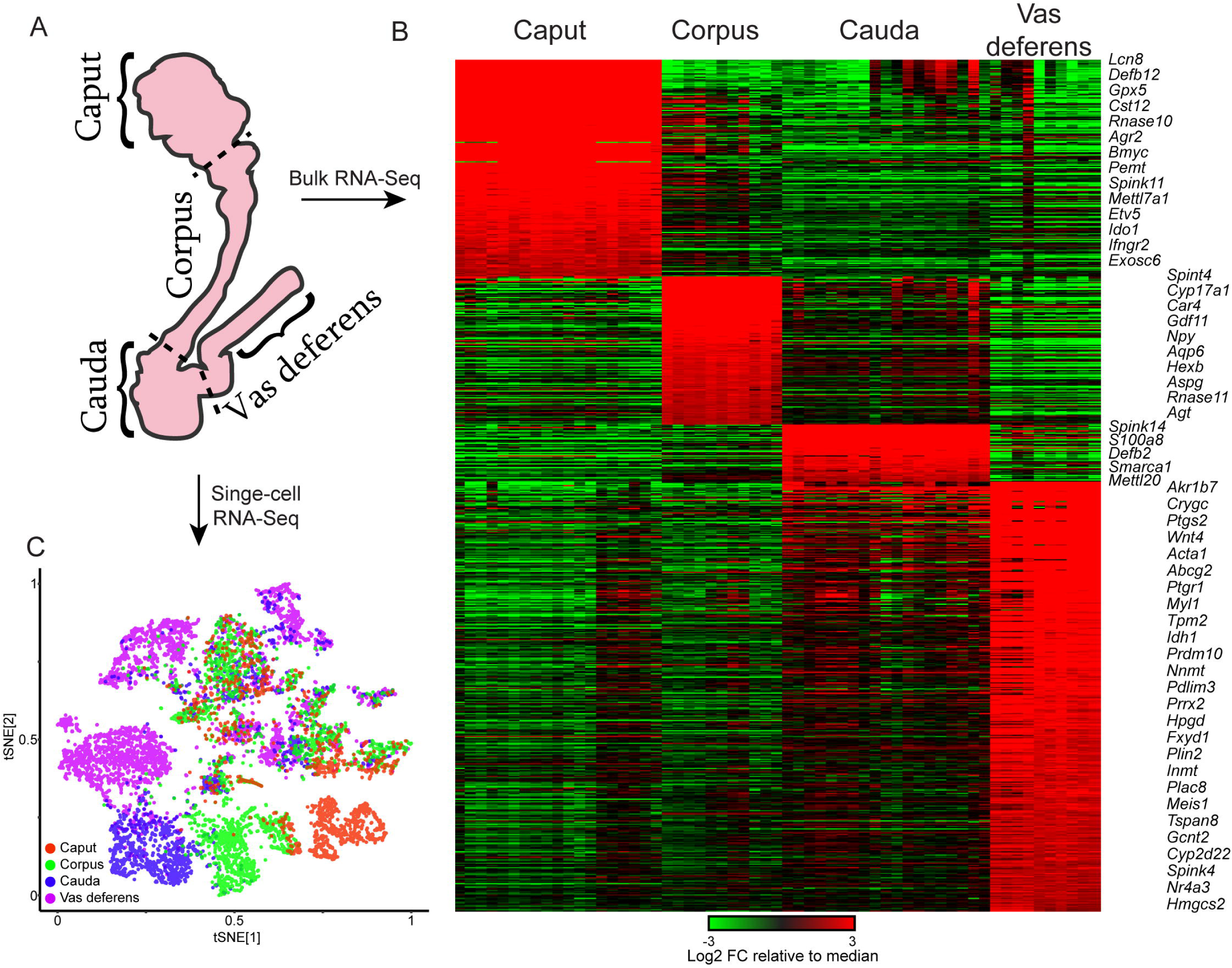
Overview of dataset. A) Cartoon showing the regions investigated here. Four anatomical regions – the caput, corpus, and cauda epididymis, as well as the vas deferens – were characterized by bulk RNA-Seq and single-cell RNA-Seq. B) Bulk RNA-Seq dataset, showing region-enriched genes (log2 fold change relative to dataset median of at least 2, with a maximum expression in one of the four dissections of at least 20 ppm). See also **Figure1-figure supplement 1**. C) Single cell RNA-Seq dataset, clustered by t-SNE and annotated according the four anatomical regions in (A).

To disentangle the contributions of the diverse cell types comprising any given region of the mouse epididymis, we used microfluidic single cell barcoding (10X Chromium) and single-cell RNA-Seq to characterize gene expression in dissociated cells from the four anatomical samples above – caput, corpus, cauda, and vas deferens. Each population represented a pool of 8 tissue samples obtained from 4 male mice sacrificed at 10-12 weeks of age. We note that dissection margins were varied slightly between individual samples to ensure complete recovery of cell types present at dissection borders.

Following single-cell RNA-Seq, quality control, and removal of reads attributable to contaminating cell-free RNA (**Methods**), we obtained a final dataset with 8,880 individual cells with an average of 3110 (median = 1965) unique molecular identifiers (UMIs) per cell. We detected cells with similar transcriptional profiles through dimensionality reduction and clustering (Methods) and the resulting map was visualized using t-distributed stochastic neighbor embedding (t-SNE), with cells from each anatomic location labeled in different colors (**Figure 1C**). Consistent with the dramatic segment-specific gene expression programs well-known to distinguish the different regions of the epididymis, we find that roughly half of the clusters are composed entirely of cells from one tissue sample – the caput, corpus, cauda, or vas (**Figure 1C**). Region-specific clusters included all principal cell clusters (discussed below), as well as muscle cells from the vas deferens samples. The remaining cell clusters included cells from multiple dissections and thus represent various cell types present throughout the epididymis.

We assigned these 21 discrete clusters to known and potentially novel cell types by the expression profiles of marker genes detected through differential expression analysis (**Methods**, **Figure 2A**, **Table S2**). This readily identified several populations of principal cells and basal cells, one of clear cells, and several for interstitial cells including fibroblasts, muscle, and immune cells (**Figure 2A**). Genes enriched in each cluster are shown in **Figure 2B**, illustrating both known and novel marker genes for various epididymal cell types. Expression of several well-known marker genes, along with genes of interest in downstream analyses, are shown across all individual cells in **Figure 2C**. Overall, we find that epididymal cells are distinguished primarily by two broad molecular features. First, functionally and histologically distinct cell types such as clear cells, basal cells, and so on are distinguished by known marker genes, as for example genes encoding vacuolar ATPase subunits (*Atp6v1a*, *Atp6v0c*, etc.) are highly expressed in clear cells which are responsible for luminal acidification (Breton et al., 1998; Breton et al., 1999; Brown et al., 1992). Second, in addition to the genes that define histologically distinct cell types, a large number of genes distinguish principal cells originating in different regions along the length of the epididymal tube. This suggests that principal cells are specialized within their niches, maintaining a complex succession of luminal microenvironments across the epididymis.

**Figure 2.**
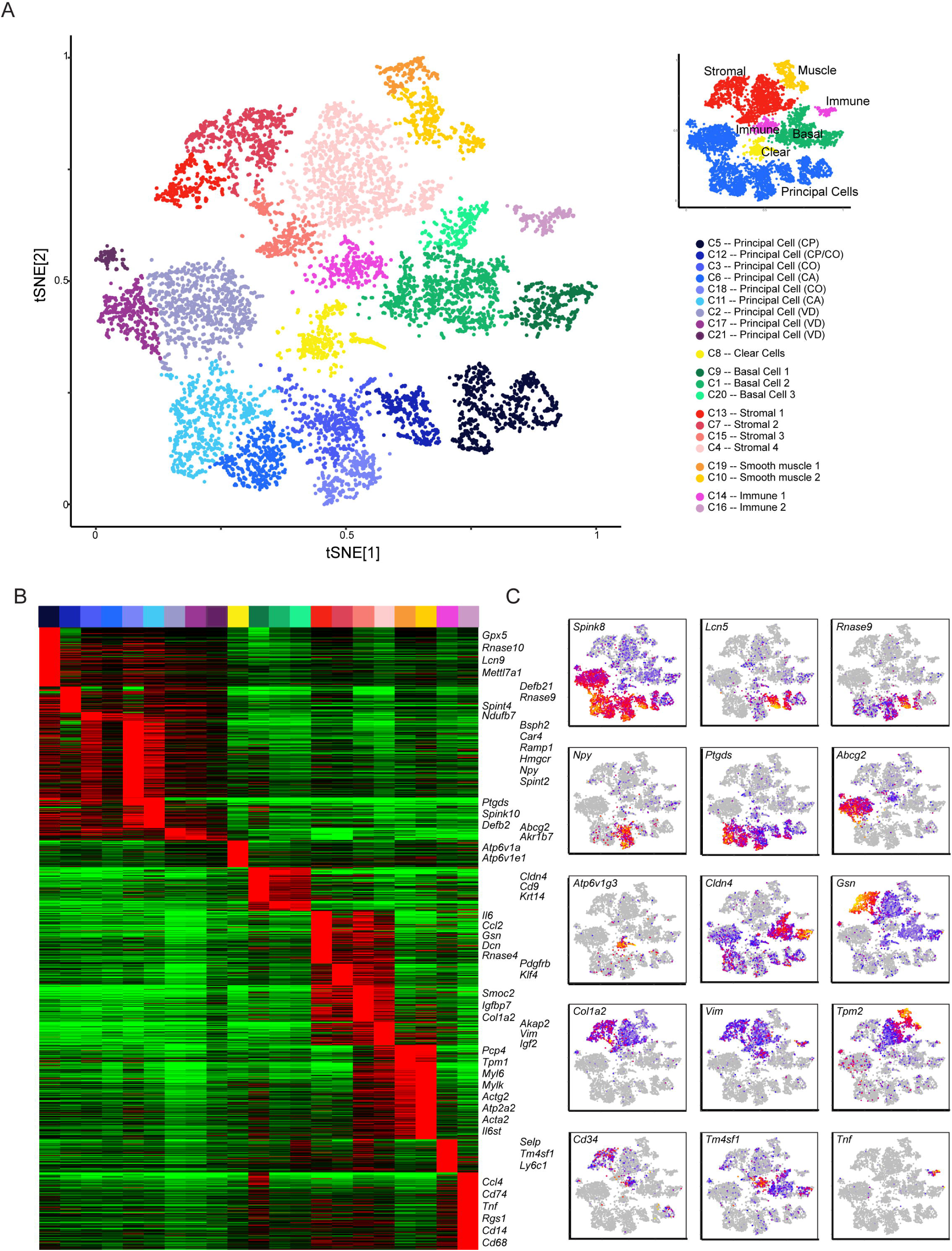
Single-cell decomposition of the epididymis. A) Single-cell cluster (expanded from **Figure 1C**) with 21 clusters annotated according to predicted cell types. Inset shows coarser cell type groupings used for downstream subclustering. B) Heatmap of genes enriched in each cluster. Among the genes exhibiting significant variation across the 21 clusters, 1241 genes enriched at least 4-fold in at least one cluster were selected for visualization. Heatmap shows these genes sorted according to the cluster where they exhibit maximal expression. C) Expression of key marker genes across the entire dataset.

Below, we provide a broad overview of the various cell populations across the epididymis, detailing key molecular features of the various cell types identified and further exploring cellular diversity.

### Segment-specific gene expression in principal cells

Principal cells are highly active secretory and absorptive cells which are responsible for producing much of the unusual protein composition of the epididymal lumen, as well as playing additional roles in modulating luminal pH, and in lumicrine signaling to other cell types. As their name suggests, principal cells comprise the major cell type present in the adult epididymis, and our data also suggest (based on shared markers – below) that the majority of vas deferens epithelial cells are analogous to epididymal principal cells. The gene expression profile of these cells allows the ready assignment of the principal cell clusters of the epididymis to anatomically-defined segments by comparison to the gold standard microarray analysis of epididymal gene expression (Johnston et al., 2005). To this set of six epididymal clusters, we added three clusters of epithelial cells from the vas deferens (**Figure 3A**).

**Figure 3.**
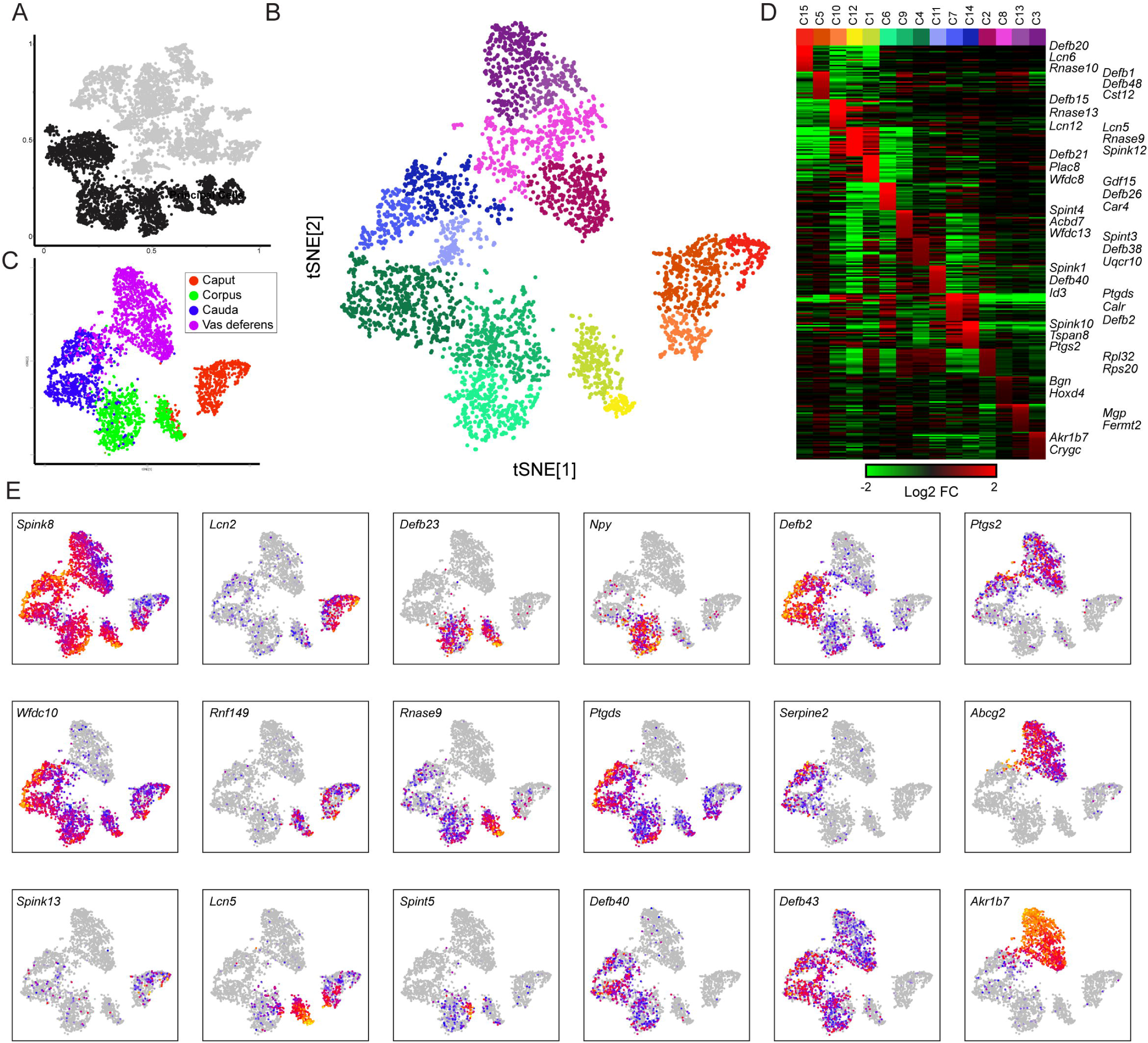
Modulation of the principal cell transcriptome across the male reproductive tract. A) Clustering from the overall dataset (**Figure 2A**), with principal cell clusters highlighted in black. B) Reclustering of extracted principal cells, visualized by t-SNE. Distinct colors highlight the 15 resulting principal cell clusters. See also **Figure 3-figure supplements 1-3**. C) Reclustered principal cells, as in panel (B), with cells colored according to anatomical origin. D) Heatmap showing the top 20 genes enriched in each of the principal cell clusters. E) Expression of key marker genes across all principal cells, visualized as in **Figure 2C**.

To investigate cellular heterogeneity among principal cells, we performed iterative re-clustering with cells from these nine clusters (**Methods**, **Figure 3B**). As expected, the second round of clustering was dominated by the anatomical origin of principal cells, separating cells from the caput, corpus, cauda, and vas deferens (**Figure 3C**). Moreover, at this resolution cells from each anatomic region could be further subdivided into up to three subtypes (**Figure 3B**), largely reflecting further subspecialization of cells across the multiple compartmentalized epididymal segments (Johnston et al., 2005). For example, within the caput epididymis, principal cells from segments 1-2 (*Lcn2*, *Cst11*) could be distinguished from cells from segments 3-4 (*Defb15*), while corpus principal cells were similarly separated into *Lcn5*/*Rnase9*-positive cells corresponding to segments 5-6, and *Npy*-positive cells corresponding to segment 7 (**Figures 3D-E**). This is consistent with prior functional and molecular studies of epididymal segmentation, and supports the notion that the anatomical segmentation of the epididymis generates multiple discrete microenvironments experienced by sperm during post-testicular maturation (Domeniconi et al., 2016).

As is clear from many prior studies of epididymal gene regulation (Guyonnet et al., 2009; Hall et al., 2007; Jelinsky et al., 2007; Jervis and Robaire, 2001; Johnston et al., 2005; Thimon et al., 2007), the genes that define principal cells from any given segment tend to be members of multi-gene families, typically encoded in genomic clusters of many paralogous genes (**Figures 3D-E, Figure 3-figure supplement 1**). Functionally, these clusters include gene families encoding serine protease inhibitors, antimicrobial defensin peptides, cysteine-rich peptides involved in glycocalyx maturation, small molecule-binding lipocalins, aldo-keto reductases, and RNaseA family members. The encoded proteins play roles throughout the male reproductive tract, both locally in the epididymal lumen, and at downstream regions of the male reproductive tract or even in the female reproductive tract. Curiously, a wide range of these region-specific proteins, including various metabolic enzymes, small molecule binding proteins, and defensins, end up decorating the sperm surface.

We next focused specifically on gene expression in the vas deferens – unlike the relatively robust literature on gene expression along the length of the epididymis, the adult vas deferens has been the subject of relatively few genome-wide studies. In contrast to the principal cells of the epididymis, few of the marker genes for the vas deferens are encoded in large multi-gene families, with two major exceptions being members of a gene family encoding serine protease inhibitors (*Akr1b7*, *Akr1c1*) (Jagoe et al., 2013) and members of crystallin-encoding genes (*Crygb*, *Crygc*). Instead, relatively highly-expressed genes in the vas deferens included various transmembrane transporters (*Abcg2*, *Fxyd4*, *Aqp2*, etc.) and signaling molecules (*Plac8*, *Ptgr1*, *Ptges*, *Ptgs2*, *Prlr*, *Cited1*, *Pcbd1*, *Comt*, *Slco2a1*, etc.). Vas deferens clusters were also enriched for expression of nuclear-encoded genes involved in mitochondrial energy production (*Atp5g1*, *Uqcr10*, *Ndufa3*, *Cox6b1*, *Ndufa7*, *Ndufa11*, *Atp5e*, *Mrpl33*, etc.), but this was not unique to the vas deferens – across the epididymis, mitochondrial energy production was overall enriched in principal cells, clear cells, and muscle cells, and depleted in basal, immune, and stromal cells (**Figure 3-figure supplement 2**).

Interestingly, several vas deferens marker genes are best-known for their roles in the placenta. For example, the xenobiotic transporter Abcg2 is localized to the apical end of syncitiotrophoblast cells of the human placenta, protecting the developing fetus by pumping small molecules back into the maternal circulation, thereby preventing xenobiotics from crossing the placental barrier (Wang et al., 2006). Abcg2 has also been implicated in sperm function, as Abcg2 is present on sperm from testicular spermatogenesis onwards, and may play a role in modulating the levels/localization of specific membrane lipids (Caballero et al., 2012). However, the function of Abcg2 in the vas deferens is unclear – interestingly, ABC-class transporters such as Abcg2 are typically localized to the apical end of polarized epithelia, which we confirmed with immunofluorescence (**Figure 3-figure supplement 3**). This suggests that Abcg2 expression in the vas deferens would be predicted to pump small molecules *into* the epididymal lumen, rather than clearing them out. As ABC class transporters such as Abcg2 are known to pump some endogenously-synthesized molecules (such as heme), our data suggest an unappreciated role for this protein in concentrating some factor(s) in the lumen of the vas deferens.

### The clear cell regulatory program

Functionally, clear cells are largely responsible for one of the best-known functions of the epididymis, which is to provide an acidic luminal environment that ensures that sperm remain quiescent until mating occurs (Breton et al., 1998; Breton et al., 1999; Brown et al., 1992). This function results from high-level expression of vacuolar ATPase (V-ATPase) genes (Breton et al., 1996; Carr and Acott, 1984), and clear cells are readily identified in our dataset by expression of the V-ATPase-encoding marker genes *Atp6v0c*, *Atpv1e1*, and *Atp6v1a* (**Figure 2C**). In addition to expressing V-ATPase subunits and genes encoding V-ATPase-interacting proteins (*Dmxl1*), clear cells were enriched for genes involved in a wide range of functions, with the two most prominent being energy metabolism and membrane trafficking. In the former case, as noted above for the vas deferens, clear cells also express high levels of genes involved in metabolic energy production, presumably serving to provide high levels of ATP needed to power the acidification of the epididymal lumen. In addition to this elevated energy production, clear cells also appear to be characterized by active membrane recycling/endocytosis, with high expression of various membrane trafficking genes (*Dab2*, *Ap1s3*, *Arf3*, *Stx7*, *Cdc42se2*). Elevated membrane trafficking activity may be related to the active intracellular trafficking required for the localization of the vacuolar ATPase to clear cell microvilli (Shum et al., 2011).

To explore potential functional diversity among epididymal clear cells, we isolated and reclustered them as described for principal cells (**Figure 4A**), finding three potential clear cell subclusters (**Figure 4B**). Two of these clusters included clear cells from throughout the epididymis, while Cluster 2 was comprised entirely of clear cells from the caput and corpus epididymis (**Figure 4C**). Prior histological studies defined a morphologically distinctive type of clear cell, known as “narrow” cells, that are uniquely found in the caput epididymis (Breton et al., 2016; Sun and Flickinger, 1979, 1980). However, no known markers have been identified for this subtype of clear cells, raising the question of whether narrow cells are functionally distinct from other clear cells. We find that clear cells in Cluster 2 were distinguished by the expression of a small number of genes, with one standout marker gene: *Iapp*, encoding islet amyloid polypeptide, or Amylin (**Figure 4D**). To validate these potentially-distinct clear cell types, we stained epididymal tissue sections for a general clear cell marker (vacuolar ATPase) and for Amylin. Consistent with the single-cell expression profiles, we find that Amylin stains only a subset of clear cells (**Figure 4E**), demonstrating the presence of two sub-types of clear cells. Notably, Amylin-positive cells were detected primarily in the caput epididymis, and to a lesser extent in the corpus, but were undetectable in the cauda or vas deferens. Amylin may therefore represent a previously-unknown marker for the so-called narrow cells, which have been described as a clear cell subtype specific to the caput epididymis. It will be interesting in future studies to determine whether Amylin-positive clear cells carry out any functions distinct from the bulk of clear cells.

**Figure 4.**
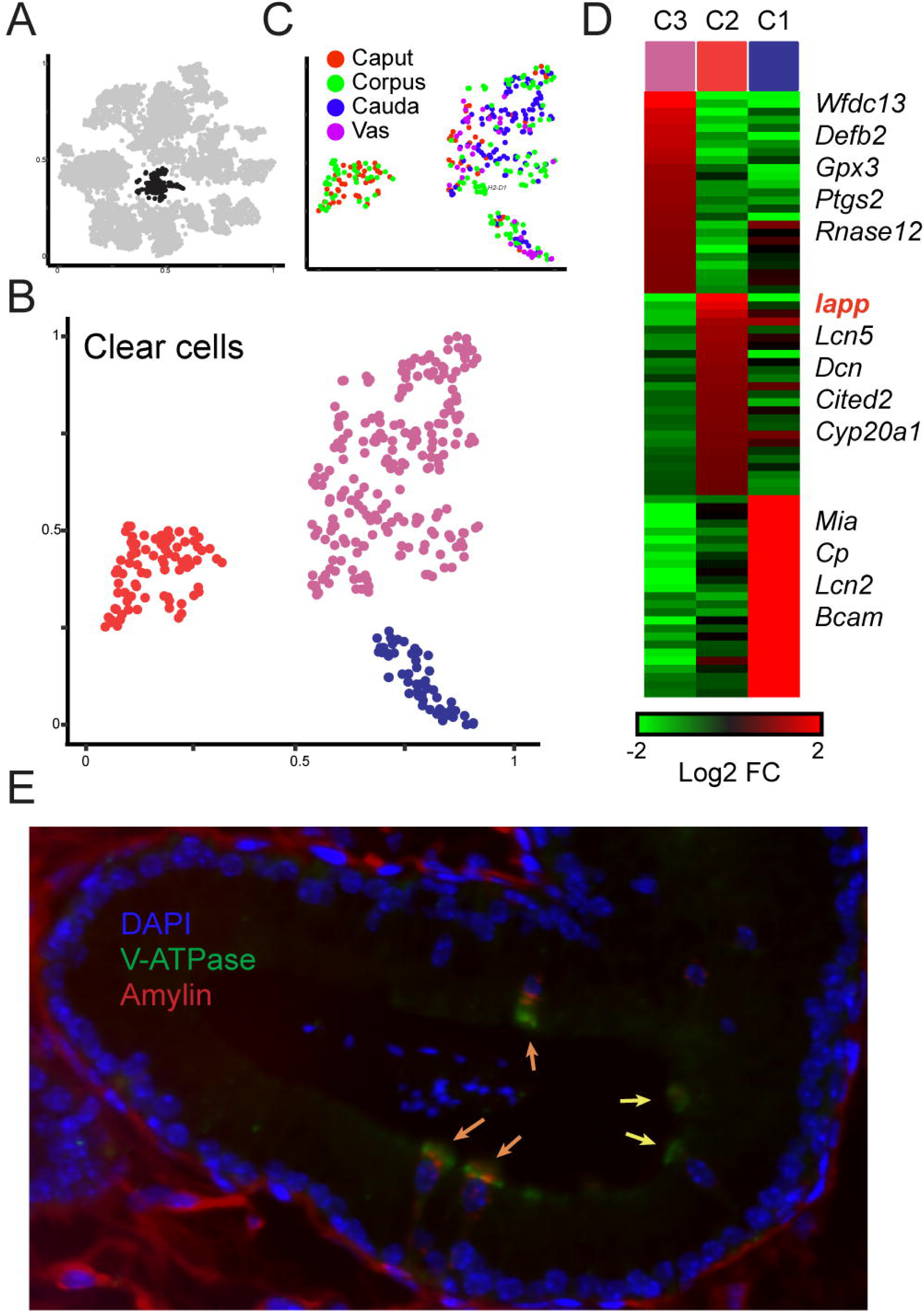
Amylin is a marker of a distinctive subset of clear cells. A-C) Reclustering of clear cells. Panels are analogous to **Figures 3A-C**. D) Heatmap showing marker genes for the three clear cell clusters. E) Amylin marks a subset of epididymal clear cells. Immunofluorescence image of a tubule of the caput epididymis, stained for DAPI, V-ATPase-G3 (marker for clear cells), and Amylin, as indicated. Yellow arrows show Amylin-negative clear cells, while orange arrows show Amylin-positive clear cells. Note that interstitial staining for Amylin is artifactual, as it was also observed in slides stained with secondary antibody alone (not shown).

### Basal cells

Another major epididymal cell type consists of the basal cells, so-named for the fact that their cell bodies are primarily localized on the basal surface of the tube, although cytoplasmic processes have been described to reach the apical surface of the epithelial sheet in some regions of the epididymis (Shum et al., 2008). Although the functions of basal cells remain somewhat obscure, they are believed to play an important role in maintaining the structural integrity of the blood-epididymis barrier, and it has recently been proposed that they may be adult stem cells for the epididymal epithelium (Mandon et al., 2015). Three clusters of basal cells were identified in our overall dataset based on the expression of marker genes including *Itga6* and *Krt14* (**Figures 2C-D**). Co-expressed with these genes were a wide variety of genes involved in cell adhesion (*Bcam*, *Gja1*, *Cldn1*, *Cldn4*, *Epcam*, *Sfn*, *Cdh16*, *Itgb6*) and intercellular signaling (*Egfr*, *Hbegf*, *Nrg1*, *Ctgf*, *Dll1*, *Egfl6*, *Adm*, *Cd44*, *Cd9*, *Tacstd2*), consistent with a recent genome-wide analysis of isolated rat basal cells (Mandon et al., 2015). In addition to expression of cell adhesion molecules, basal cells exhibited high level expression of several genes involved in membrane trafficking and lipid metabolism (*Cd9*, *Sdc1*, *Sdc4*, *Vmp1*, *Apoe*, *Apoc1*, *Sgms2*), suggesting that basal cells may exhibit more dynamic membrane remodeling, or vesicle production, than currently appreciated.

To further explore the diversity of basal cell subtypes in our dataset, we extracted and reclustered the three putative basal cell clusters as described above for principal and clear cells. After removing a small group of macrophages present in the initial clustering (**Figure 5-figure supplement 1A**), we found relatively little diversity among basal cells (**Figure 5-figure supplements 1B-D**). The primary distinction between different basal cell subclusters was based on expression of ribosomal protein genes. However, we found that these two groups (RPG high and RPG low) also exhibited significant differences in the number of UMIs captured per cell (**Figure 5-figure supplements 1D-E**). Therefore, it is unclear whether the RPG-low basal cells represent a biologically-meaningful population marked by low biosynthetic activity, or whether these cells represent technical artifacts (of, say, inefficient lysis and RNA recovery). Beyond this major grouping of basal cells, we did identify one additional cluster of possible interest, with a subset of basal cells expressing high levels of *Notch2*, *Spry2*, *Pecam1*, *Slc39a1*, *Ccl7*, *Cd93*, and *Penk*. These basal cells may potentially represent an interesting topic for future followup studies.

### Supporting cell types: stromal, immune, and muscle cells

In addition to the major cell types that comprise the lining of the tube and shape sperm maturation, the epididymis is surrounded by an array of supporting cell types including stromal cells and infiltrating immune cells. In addition, the distal epididymis – the cauda epididymis and in particular the vas deferens – is surrounded by a muscular sheath whose contraction drives sperm movement during ejaculation. Eight of the clusters in our initial clustering analysis clearly represent these various supporting cell types. Two clusters of muscle cells, defined by their expression of various actomyosin cytoskeletal genes (*Myh11*, *Mylk*, *Acta1*, *Tpm2*, etc.) are clearly identified, and as expected are primarily populated by cells from the vas deferens dissection (**Figures 1C, 2**). Muscle markers are also expressed at lower levels in a subset of the collagen-enriched stromal cell clusters (**Figure 2C**), consistent with the potential presence of myofibroblasts (see below). Other supporting cell types included endothelial cells, fibroblasts, macrophages, and other immune cells.

To further explore the cellular diversity among the supporting cell types, we isolated the cells from the three sets of clusters corresponding to muscle cells, stromal cells, and immune cells, and subjected each group of cells to a second iteration of clustering (**Figure 5, Figure 5-figure supplement figure 2**). We found relatively little variation among the muscle cells, which separated into two clusters (**Figure 5-figure supplement figure 2**). These clusters were distinguished primarily by the expression levels of ribosomal protein genes, similar to the major distinction between basal cells with high and low UMI coverage (**Figure 5-figure supplement figure 1**). Although in the case of these muscle cell subclusters we did not observe substantially different UMI coverage (not shown), nonetheless we were not confident that these are biologically-relevant subclusters, and so muscle cells were not further evaluated.

**Figure 5.**
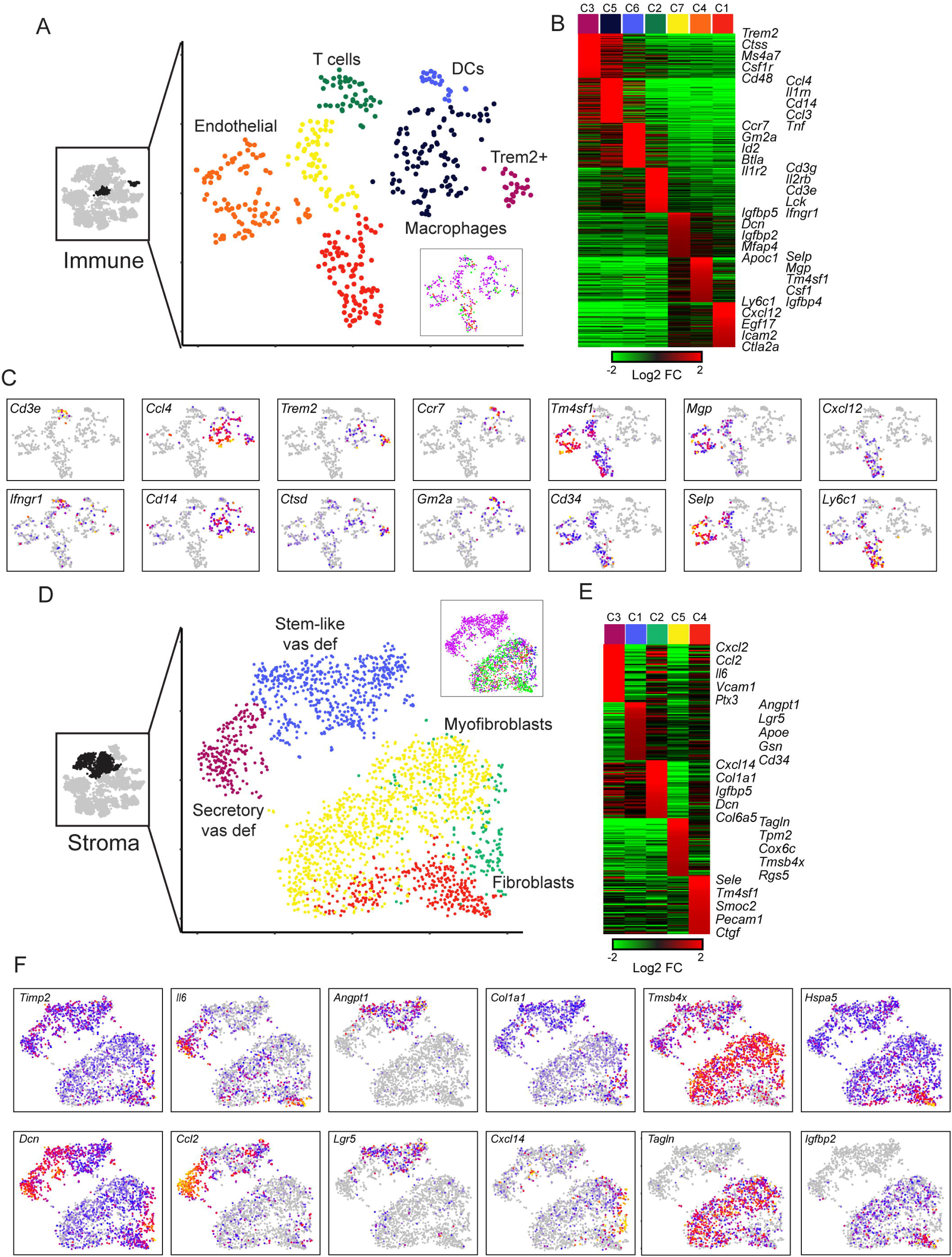
Diverse stromal cell populations in the epididymis. A) Reclustering of immune and endothelial cells. Left panel highlights the clusters from full dataset used for reclustering, while right panels show reclustered data. Inset shows cells colored according to the anatomical region where they were found. B) Heatmap of the top 100 markers for each of the immune subclusters in (A). C) Expression of individual marker genes across immune and endothelial cell subpopulations, as in **Figure 2C**. D-F) Reclustering of stromal cells, arranged as in panels (A-C). Heatmap (E) shows the top 50 genes for each subcluster.

Turning next to immune cells (**Figures 5A-C**), subclustering of the two initial clusters revealed a range of infiltrating immune cell populations, as well as endothelial cells, including:

- Macrophages expressing *Ccl3*, *Cll4*, *Cd14*, *Il1rn*, *Ifnb1*, *Tnf*, etc.
- A distinct subcluster of potentially anti-inflammatory Macrophages, expressing high levels of *Trem2* and *Wfdc17*, as well as *Ctsd*, *Dcxr*, *Ctsa*, *Ctsb*, *Csf1r*, *Rnase4*, and *Lyz2*.
- Dendritic cells expressing *Ccr7*, *Kit*, *Gm2a*, *Btla*, and various MHC genes (*H2-0a*, *H2-Aa*, *H2-Ab1*, *H2-DMa*).
- T cells, expressing *Cd3e*, *Cd3g*, *Cd3d*, *Il2rb*, *Lck*, *Cd7*, and *Gzmb*.
- Endothelial cells, expressing *Cd34*, *Tm4sf1*, *Icam2*, and *Emcn*. Interestingly, endothelial cells separated into three subclusters, with the two most distinctive clusters (C1 and C4) distinguished by different intercellular signaling and adhesion molecules (C1: *Cxcl12*, *Egf17*, *Dll4*, *Ly6a*, *Ly6c1*; C4: *Selp*, *Sele*, *Mgp*, *Csf3*, *Csf1*, *Vcam1*, *Gsn*).

Among these cell types, the identification of macrophages and T cells is notable as macrophages and infiltrating lymphocytes represent the predominant groups of immune cells involved in the blood-epididymis barrier (Da Silva and Barton, 2016; Serre and Robaire, 1999). These cells correspond to the infiltrating immune populations identified in early histology studies as somewhat poorly-defined “Halo” cells (Serre and Robaire, 1999; Sun and Flickinger, 1979). Interestingly, we found a strong bias for macrophages to be found in the vas deferens dissection (p<10^-7^, hypergeometric), suggesting two possibilities. One is that the close association of the vas deferens with blood vessels could simply result in more circulating macrophages being captured in this dissection. Alternatively, we find some evidence (discussed below) that vas deferens stromal cells may play an active role in recruiting macrophages.

We finally turned to the four clusters from the full dataset putatively annotated as stromal cells, based on their relatively common expression of various collagen genes (**Figure 2B**). After reclustering (**Figure 5D-F**), we find that the largest subcluster of these cells was characterized by high levels of muscle markers (albeit not as high as the bona fide muscle cells – **Figure 2C**) such as *Tagln*, *Myl9*, *Tpm2*, *Acta2*, etc., and thus presumably correspond to the myoid cells whose contraction plays roles in moving sperm and fluid through the epididymal lumen (Oliveira et al., 2016). Two other clusters of fibroblasts (C2, C4) were distinguished by relatively few marker genes and are not further considered. Finally, we identified two particularly interesting fibroblast subtypes that were almost entirely confined to the vas deferens (**Figure 5D**, inset). One of these clusters expressed a number of markers associated with stem cells (*Angpt1*, *Lgr5*, *Tspan8*) and may represent mesenchymal stem cells. The other vas-specific cluster was distinguished by high level expression of a large number of intercellular signaling molecules (*Il6*, *Cxcl1*, *Cxcl2*, *Cxcl12*, *Ccl2*, *Ccl7*, *Vcam1*, *Igf1*, etc.) and so were tentatively annotated as “secretory fibroblasts.” It will be interesting in future studies to investigate the biology and functions of these unusual cell populations in the vas deferens.

### Cell to cell communication pathways in the epididymis

We noted that many of the cell subpopulations defined throughout this study were distinguished by expression of intercellular signaling molecules, from adhesion molecules to secreted cytokines and their receptors. Prior studies decomposing complex tissues into single-cell RNA-Seq have leveraged the expression profiles for known ligand-receptor pairs to infer significant pathways for cell to cell communication (Cohen et al., 2018). We therefore turn finally to exploration of potential intercellular communication networks throughout the epididymis.

We separately identified known ligands, and their receptors, expressed across 34 cell subpopulations defined in this study (**Table S3**), using a large-scale resource (Ramilowski et al., 2015). From these lists, we identified the number of potential interactions linking every cell type to every other cell type (**Figure 6**). Of these, significant interactions between cell types were defined based on the number of cases of expression of a ligand-receptor pair between the cell types, relative to the number of pairs expected by chance based on the total number of ligands and receptors expressed in each cell type (**Methods**). This revealed multiple potential pairs of cells in communication, of which we highlight two. First, we find that basal cells and endothelial cells expressed complementary sets of ligands and receptors, consistent with the location of basal cells at the periphery of the epididymal epithelium. Second, we found reciprocal connections between macrophages and the “secretory fibroblasts” in the vas deferens, suggesting the possibility that these vas deferens stromal cells play a role in recruiting macrophages to this tissue.

**Figure 6.**
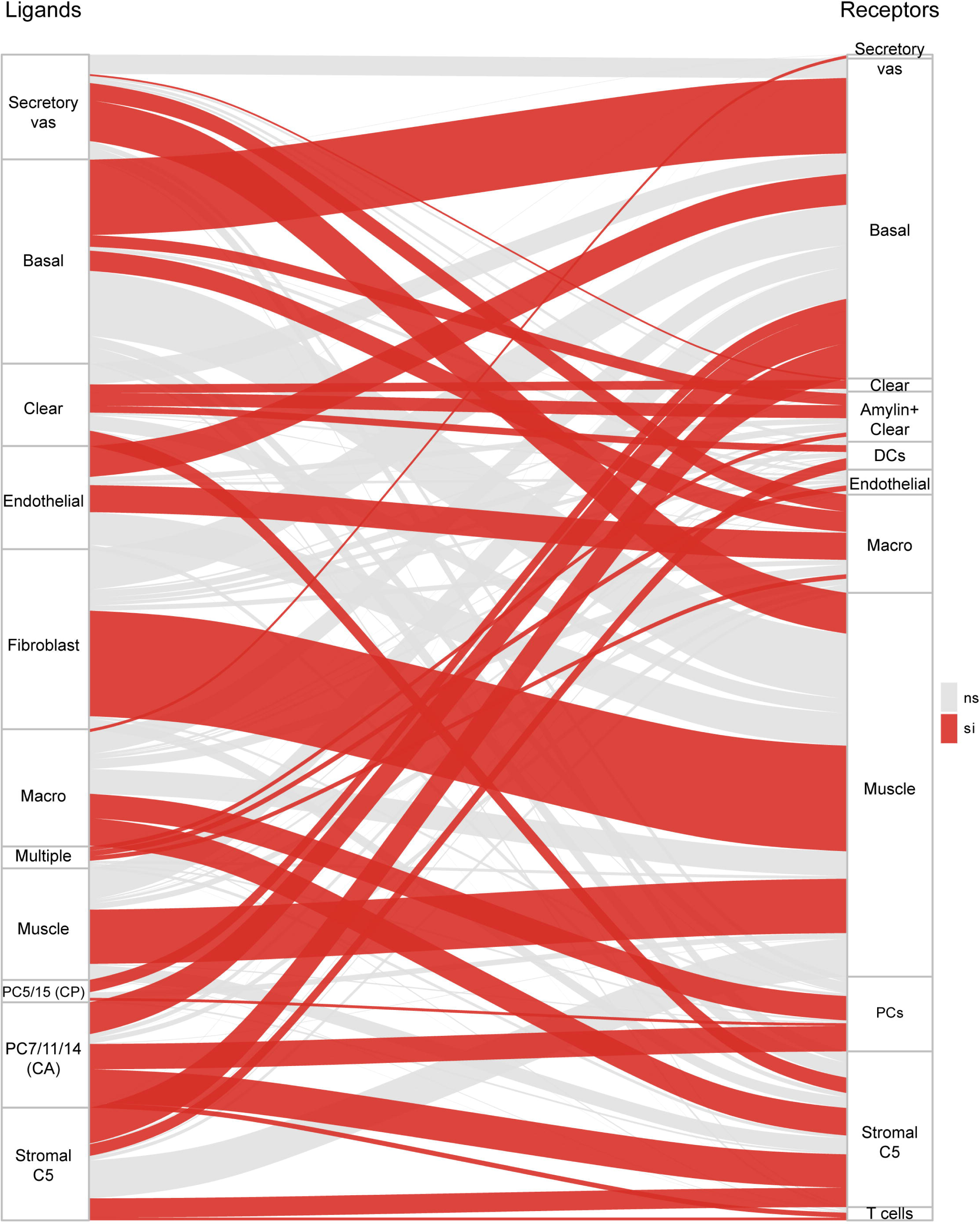
Potential signaling interactions between epididymal cell types. Bars on left and right show clusters of ligands and receptors (see **Table S3**) named according to the cell population with highest expression levels. Rectangle lengths represent the number of putative connections between ligands (left panels) and receptors (right panels) expressed in different cell types. Lines connecting rectangles represent connections between matching ligand-receptor pairs expressed. If the number of interactions between a cell type is equal or less than the number expected by chance (based on the number of expressed ligands and receptors in the pair of cell types, hypergeometric test), lines are grey, while lines are colored red for cell type pairs with a significant enrichment of matching ligand-receptor pairs. See **Table S3** for ligand and receptor expression in the 34 cell populations.

## DISCUSSION

Here, we present a single-cell atlas of the murine epididymis and vas deferens, building on prior histological and marker-based surveys of the epididymis. We recover all previously-described cell types in the murine epididymis, including support cells which have not been characterized in detail in this tissue, and provide more complete molecular profiles for all of these cell types. Our data provide a compendium of transcriptional profiles for a range of cell types in this key reproductive organ, identify markers for potentially novel cell types in this tissue, and motivate a wide variety of new hypotheses for future molecular followup studies.

### Distinctive principal cell programs along the length of the epididymis

The most striking feature of the epididymis is the presence of highly distinctive regional gene expression programs, manifest primarily in the principal cells that comprise the majority of the epithelial cells of the tube. Consistent with many previous surveys of the epididymis in a variety of mammals, we find multiple discrete groups of principal cells arrayed along the length of the epididymis, with roughly eight unique gene expression signatures being appreciable across the epididymis and vas deferens (**Figure 3D**). The genes enriched in specific regional domains are involved in a wide range of functions, and include signaling peptides (*Npy*), putative RNA modifying enzymes (*Mettl7a1*), and metabolic enzymes (*Idi1*, *Fdps*). However, the most notable feature of region-specific gene regulation in the epididymis is the highly segmental expression of individual members of several large multi-gene families that are often organized in genomic clusters, including β-defensins, aldo-keto reductases, lipocalins, serine protease inhibitors, and RNases (**Figure 3-figure supplement 1**).

The activities encoded by these gene clusters encompass many of the best-characterized functions of the epididymis, as for example glycocalyx and membrane remodeling of the sperm surface is one of the first processes identified to occur in this tissue (Tecle and Gagneux, 2015), while the wide array of antimicrobial peptides presumably serve to insulate the testis from ascending infections (Gregory and Cyr, 2014; Hall et al., 2007). Among the other gene families, aldo-keto reductases are involved in the biosynthesis of signaling molecules (Volat et al., 2012), including prostaglandins (Kabututu et al., 2009), which have the potential both to mediate lumicrine signaling within the male reproductive tract as well as signaling between partners following delivery to the female reproductive tract. The aldo-keto reductases may also protect sperm from oxidative damage during transfer outside the body. The lipocalins are a family of small molecule binding proteins which bind to lipophilic cargoes with a range of functions from signaling (retinoic acid, (Lareyre et al., 1998; Ong et al., 2000)) to innate immunity (siderophores, (Golonka et al., 2019)). Serine protease inhibitors are homologous to the semen coagulum proteins (which are synthesized primarily by the seminal vesicle) and potentially play some role in the protease-controlled processes of semen coagulation and liquefaction (Clauss et al., 2011), but may also be involved in acrosome maturation and sperm capacitation (Ma et al., 2013). Finally, high level expression of various RNases presumably explains the extraordinarily high levels of tRNA cleavage in this tissue (Sharma et al., 2016), with resulting tRNA fragments potentially acting as another layer of protection from selfish elements (Martinez et al., 2017; Schorn et al., 2017), or serving as environmentally-modulated signaling molecules delivered from sperm to the zygote upon fertilization (Chen et al., 2016; Rompala et al., 2018; Sharma et al., 2016).

The organization of epididymal gene expression in this way raises two obvious questions. The first is the mechanism by which individual members of these gene families are specifically expressed in individual spatial domains – what transcription factor or post-transcriptional regulator (Bjorkgren et al., 2012) is responsible for, say, expression of *Defb21* but not *Defb2* in the caput epididymis? We note that there does not appear to be any clear logic relating spatial expression with genomic location within a cluster, as most famously seen in the Hox clusters. The second question is what biological function is served by this staged expression program, which results in sperm transiting consecutive microenvironments each bearing individual members of these various gene families. In other words, is there any utility to having sperm first incubate for days in the presence of Defb21, then move into an incubation with Defb48, etc.? Would sperm function any differently if the defensins (or Rnases, etc.) expressed switched positions?

### A novel subclass of clear cells

A common goal of single-cell RNA-Seq dissection of multicellular tissues is to identify novel cell types, particularly rare cell types that might have been missed in classic histological studies. Here, we identified several cases in which we could distinguish several previously-unknown subgroups of well-known epididymal cell types. We followed up on a subdivision among clear cells, where a subgroup of clear cells confined to the caput and corpus epididymis could be distinguished by expression of *Iapp* (**Figure 4**). We confirmed the presence of both Amylin-negative and Amylin-positive clear cells, and found that these subgroups could even be found cohabitating within the same region of the epididymis. We speculate that the Amylin-positive clear cells correspond to the classic description of narrow cells in the caput epididymis (Sun and Flickinger, 1979, 1980), for which molecular markers had not previously been identified. That said, narrow cells have generally been described as confined to the initial segment of the epididymis, while we find Amylin-positive clear cells in both caput and corpus epididymis. Whatever the correspondence to narrow cells, it will be important in future studies to determine whether functional differences can be identified that distinguish Amylin-positive and -negative clear cells – Amylin has been implicated in systemic control of metabolism (Boyle and Le Foll, 2019), so it will be interesting to determine whether clear cell-secreted Amylin is used to signal locally within the male reproductive tract, or whether systemic release of Amylin is used to modulate organismal metabolism potentially in response to reproductive functionality.

### Perspective

Together, our data provide a bird’s eye view of the cellular composition of the murine epididymis. We identified several novel features of epididymal cell biology, most notably including a detailed examination of the understudied vas deferens. Potentially-novel cell types identified include Amylin-positive clear cells, as well as two subtypes of fibroblasts − secretory and stem-like – confined to the vas deferens. Moreover, our data highlight several groups of cells with the potential for extensive intercellular signaling, and suggest that secretory fibroblasts in the vas deferens may play roles in recruiting macrophages. These data will provide a rich resource for hypothesis generation and future efforts to more fully understand the biology of the epididymis.

## ACKNOWLEDGEMENTS

We would like to thank the Luban and Greer labs for providing instruments and guidance with single cell sequencing, and members of the Rando and Garber labs for critical reading of the manuscript and insightful discussions. This work was supported by NIH grant R01HD080224 and Templeton Foundation grant 61350. US is supported by NIH grant 1DP2AG066622-01.

## METHODS

### Mice

Tissues were obtained from 10-12 week old male FVB/NJ mice. All animal care and use procedures were in accordance with guidelines of the University of Massachusetts Medical School Institutional Animal Care and Use Committee.

### Dissection

In order to obtain a sperm-depleted single cell suspension, eight epididymides from four 10-12 weeks old FVB mice, euthanized according to IACUC protocol, were dissected into a 10cm plate containing 10ml of Krebs media pre-warmed to 35C. Using a dissection microscope with warm stage set to 35C, the organ was cleared from any fat and excessive connective tissue. In a clean 10cm dish containing 10 ml of Krebs media the organ was cut into four segments that roughly corresponded to caput, corpus, cauda, and vas. Each group of eight segment was placed into different 5 cm petri dishes with about 2 ml of Krebs media. Using curved scissors each group was further cut into roughly one to five-millimeter pieces and transferred to a 15ml conical tube. Media was added to a final volume of 10 ml, and samples allowed to decant while in an upright position at a 35C incubator for three to five minutes. Supernatant was discarded and this wash step repeated for two more times. In order to further remove sperm, the tubules were transferred to a 50ml conical tube containing 25 ml pre-warmed RPMI-1640 media. After ten minutes incubation at 35C under mild agitation, sample was allowed to settle for three to five minutes and supernatant discarded. These steps were repeated until supernatant looked clear.

### Dissociation

Tissue dissociation was optimized to obtain a single cell suspension free of sperm. Once dissected samples were cleared of any visible sperm, 3 ml of the RPMI-1640 sample mixture was transferred to a small 25ml glass Erlenmeyer flask containing 7 ml of dissociation media (Collagenase IV 4mg/ml, DNAse I 0.05 mg/ml) and placed in a 35C water bath with rotations of 200 rpm for 30 to 45 minutes. Afterwards, samples were transferred to a conical tube and allowed to settle. Supernatant was removed, leaving four to five ml of sample solution to be returned to the Erlenmeyer flask, were 10 ml of 0.25% trypsin-EDTA and 0.05 mg/ml DNAse and returned to the water bath. After 35 minutes the samples were pipetted up and down until there were no observable pieces of tissue, and allowed to incubate in the water bath for 20 more minutes. Once cell disaggregation was achieved 1 ml of FBS was added to inactivate the trypsin. Sample was transferred to falcon tube by passing the cell solution through a series of 100um, 70um, and 40um cell strainers, and adding media (RPMI-1640 supplemented with 10% FBS and antibiotic and anti-mycotic - 100 units/mL of penicillin, 100 µg/mL of streptomycin, and 0.025 µg/mL of Amphotericin B) to a final volume of 30 ml. Centrifuge for 15 minutes at 400 rcf (g) at RT. Discard supernatant and add 25 ml media and centrifuge for 5 minutes at 800 rcf (g) at RT. Discard supernatant and add 2ml of PBS, transfer cells to a 2 ml microfuge tube, centrifuge at 900rcf (g) for 3 minutes, discard supernatant, resuspend in 600ul PBS and count cells (tryphan blue) put them on ice and if good (>85% survival) proceed with the 10X protocol/dilutions.

### Library protocol

Single-cell sequencing libraries were prepared using Chromium Single Cell 3’ Reagent Kit V2 (10X Genomics), as per the manual, and sequenced on HiSeq 4000 at the UMass Medical School Deep Sequencing Core.

### Data processing

Sequencing reads were processed as previously described (Derr et al., 2016), using the graphical interface DolphinNext [**doi:** https://doi.org/10.1101/689539]. Scripts and the full pipeline can be accessed through GitHub (<u>github.com/garber-lab/inDrop_Processing</u>). Briefly, fastq files were generated using bcl2fastq and default parameters. Valid reads were extracted using umitools [doi:10.1101/gr.209601.116] and valid barcode indices supplied by 10X Genomics. Reads where the UMI contains a base assigned as N were discarded. Remaining reads were aligned to the mm10 genome using HISAT2 v2.0.4 (Kim et al., 2019) with default parameters and the reference transcriptome RefSeq v69. Alignment files were filtered to contain only reads from cell barcodes with at least 100 aligned reads and were submitted through ESAT (github.com/garber-lab/ESAT) for gene-level quantification of unique molecule identifiers (UMIs) with parameters -wLen 100 -wOlap 50 -wExt 1000 -sigTest .01 - multimap ignore -scPrep. Finally, we identify and correct UMIs that are likely a result of sequencing errors, by merging the UMIs observed only once that display hamming distance of 1 from a UMI detected by two or more aligned reads.

Barcodes with at least 350 and no more than 15,000 unique molecular identifiers (UMIs) mapped to known genes were considered valid cells. Additionaly, background RNA contamination was removed from the gene expression matrix using a modified version of the algorithm SoupX v0.3.1 [https://doi.org/10.1101/303727]. Briefly, barcodes corresponding to empty droplets were selected per sample according to their distribution of UMIs per barcode (caput: 20=<x=<175, corpus: 20=<x=<150, cauda: 20=<x=<150, vas: 20=<x=<115; where x=total UMIs for a given barcode). UMI counts for empty droplets were used as input for the SoupX algorithm to infer the expression profile from the ambient contamination per sample. The main alteration consisted in the selection of the fraction of contamination per cell (rho), which was determined by the most frequent number of UMIs for the empty droplets in each sample (caput=139; corpus=114; cauda=109; vas=84), under the assumption that the same contamination level could be expected from valid cell containing droplets. Using these estimated contamination fractions, the SoupX algorithm was used to remove counts from genes that had a high probability of resulting from ambient RNA contamination for each cell with the function adjustCounts.

### Dimensionality reduction and clustering

Gene expression matrices for all samples were loaded into R (V3.6.0) and merged. The functions used in our custom analysis pipeline are available through an R package (github.com/garber-lab/SignallingSingleCell). Using the clean raw expression matrix, the top 20% of genes with the highest coefficient of variation were selected for dimensionality reduction. Dimensionality reduction was performed in two steps, first with a PCA using the most variable genes and the R package irlba v2.3.3, then using the first 7 PCs (>85% of the variance explained) as input to tSNE (Rtsne v0.15) with parameters perplexity = 30, check_duplicates = F, pca = F (van der Maaten and Hinton, 2008). Clusters were defined on the resulting tSNE 2D embedding, using the density peak algorithm (Rodriguez and Laio, 2014) implemented in densityClust v0.3 and selecting the top 30 cluster centers based on the □ value distribution (□=□ × □; □ = local density; □= distance from points of higher density). Using known cell type markers, clusters were used to define 6 broad populations (Immune, Stromal, Basal, Principal, Clear and Muscle). UMI counts were normalized using the function computeSumFactors from the package scran v1.12.1 (Lun et al., 2016), and parameter min.mean was set to select only the top 20% expressed genes to estimate size factors, using the 6 broad populations defined above as input to the parameter clusters. After normalization, only cells with size factors that differed from the mean by less than one order of magnitude were kept for further analysis (0.1x(□(□/N)) > □ > 10x(□(□/N)); □= cell size factor; N = number of cells). Normalized UMI counts were converted to gene fractions per cell and the resulting matrix was used in a second round of dimensionality reduction and clustering. The top 15% variable genes were selected as described above and used as input to PCA and the first 24 PCs that explained >95% of the variance were used as input to tSNE. The resulting 2D embedding was used to determine 21 clusters as described above. Marker genes for each cluster were identified by a differential expression analysis between each cluster and all other cells, using edgeR (McCarthy et al., 2012), with size factors estimated by scran and including the sample of origin as the batch information in the design model. Each of the 6 broad cell type populations was independently re-clustered following the same procedure described above, to reveal more specific cell types. Expression values per cluster were calculated by aggregating gene expression values from individual cells in each cluster and normalizing by the total number of UMIs detected for all cells in that cluster, multiplied by 10^6^ (UMIs per million).

An R script for the steps described here and containing all parameters used is available through GitHub (<u>github.com/elisadonnard/SCepididymis</u>).

### Receptor-ligand network analysis

Receptors and ligands expressed in the dataset were selected based on a published database (Ramilowski et al., 2015). The database genes were converted to their respective mouse homologs using the Homologene release 68. Genes were filtered to select only those expressed by at least 20% of cells in at least one cell population, resulting in a set of 226 ligands and corresponding 212 receptors. Aggregated expression values were calculated based on this filtered gene matrix per cell type as described above. The annotation of each pair of expressed ligands and receptors in the data was done using the function *id_rl* from the package *SignallingSingleCell*. The expression matrix for all annotated ligands or receptors was clustered using *k-means* (k=13 ligands; k=14 receptors) and clusters were labeled and merged based on the cell population showing highest expression of those genes, resulting in 11 final ligand cell groups and 11 final receptor cell groups (**Table S3**). The frequency of connections from each ligand to each receptor cluster was calculated and significant cell-cell interactions (p<0.01) were determined with a hypergeometric test (**Figure 6**).

### Histology

Male mice from the same strain and age as previously described were anesthetized and perfused with a solution of phosphate-buffered saline (PBS) followed by 4% paraformaldehyde (PFA)/PBS, and epididymides explanted and further incubated in fixative at 4C overnight (ON). After washing the excess of PFA with PBS, the sample was incubated at 4C with a 30% sucrose, 0.002% sodium azide (NaAz) in PBS solution for 32 hours, after which the same volume of optimal cutting temperature compound (OCT) was added to the vial and kept ON at 4 C under agitation. Samples were than mounted and frozen at −80°C until sectioning. Sectioning was done at a thickness of 5 um thickness by the UMASS morphology core. Slides were stored frozen at −20C. For Vas deferens Abcg2 staining, tissues were dissected and directly placed in 2ml of 4% PFA (freshly prepared) and incubated at 4C ON with gentle shaking. The samples were further processed as described above. Tissue sections were sectioned at 8-10 um thickness using Leica 3050 cryostat. Tissue sections were transferred to Superfrost/Plus microscope slides and stored at 80C until used for immunostaining.

### Immunofluorescence

Slides were placed at a 37C warm plate for 10 minutes to ensure proper attachment of the section to the slide, then washed three times of 5 minutes in PBS 0.02% tween 20 (PBS-T) to remove OCT. Staining was performed as suggested by the anti-body manufacturer. Briefly a 5 minutes permeabilization step in PBS 0.02% triton-X was performed prior to a 30 minutes blocking with 5% goat serum 3% BSA in PBS. The primary antibodies used were V-ATPase G3 rabitt polyclonal antibody (Invitrogen) and [R10/99] Amylin mouse monoclonal (Abcam). Slides were incubated at 4C for 48hrs with the aforementioned primary antibodies (at 1:100 dilutions), and subsequently incubated with Alexa Fluor secondary antibodies for 3 hours followed by DAPI dye for 5 min. Slides were mounted with ProLong gold Antifade (Thermofisher). For Abcg2 immunostaining, slides were placed at room temperature for 10 minutes, and rehydrated in 1XPBS for 15 minutes, followed by permeablization with 1% SDS for 4 mins and 3 washes with 1XPBS. The slides were then blocked with 1% BSA, and incubated with primary antibody anti-Abcg2 (ab24115 Rat monoclonal, Abcam) ON at 4C. The slides were then washed and incubated with secondary antibody Alexa-Flour 647 for 1 hour at RT, followed by washing and DAPI staining. The slides were mounted using Fluoromount G and imaged on Axioimager (Zeiss).

### Data availability

Data will be publicly available at GEO, Accession #XXXXXXXX.

## SUPPLEMENTARY INFORMATION

### SUPPLEMENTARY TABLES

**Table S1. Bulk RNA-Seq dataset.** RNA-Seq for caput (CP), corpus (CO), and cauda (CA) epididymis, and for vas deferens (VD). All data are normalized to parts per million.

**Table S2. Marker gene expression across 21 cell clusters.** Log2 fold enrichment for 11089 genes across the 21 cell clusters (**Figure 2A**) from the full dataset.

**Table S3. Ligand and receptor expression across epididymal cell types.** Relative expression levels for all ligands or receptors expressed (>=1 UMI) in at least 20% of cells from one or more of the 34 different cell types listed.

### SUPPLEMENTARY FIGURES

**Figure 1-figure supplement 1.**
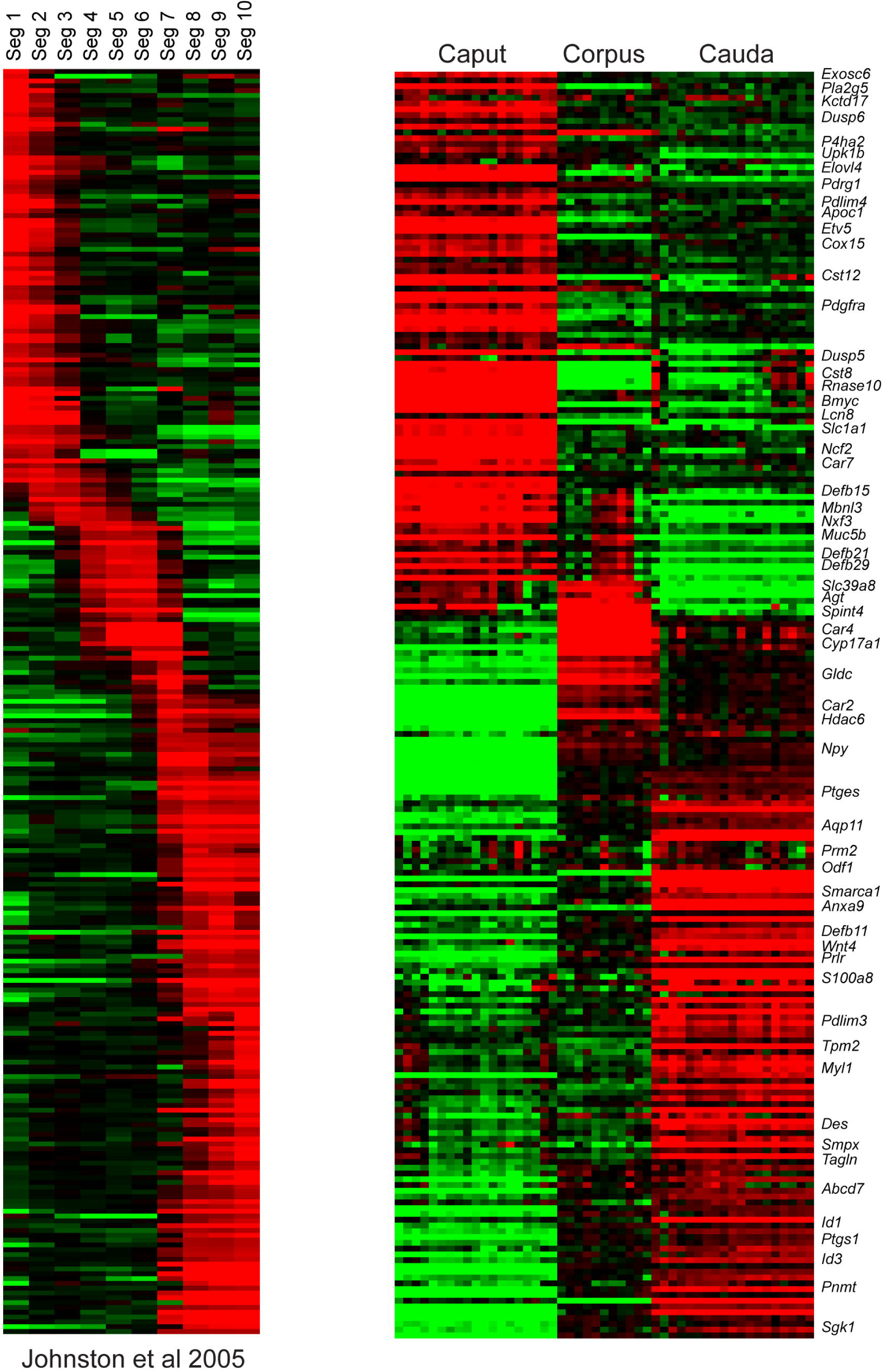
Comparison of RNA-Seq to prior microarray analysis of segmental gene expression. Left panel shows microarray data from Johnston et al, 2005 (Johnston et al., 2005), sorted according to the epididymal segment exhibiting maximal expression. Right panel shows data for the same genes in the bulk RNA-Seq data from this study, for caput, corpus, and cauda dissections.

**Figure 3-figure supplement 1.**
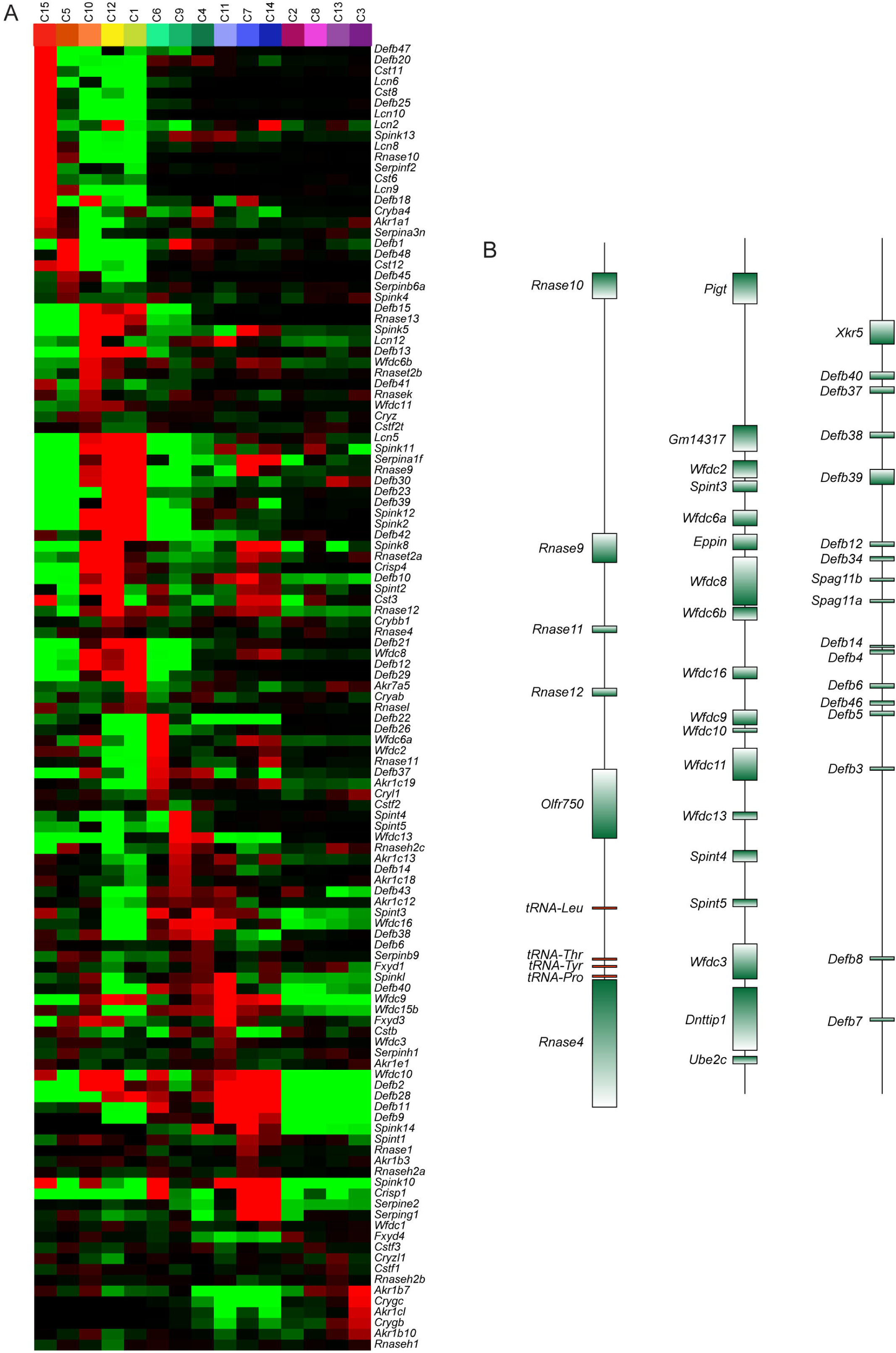
Region-specific expression of multi-gene family members. A) Heatmap showing expression of individual members of several multi-gene families (*Rnase*, *Lcn*, *Cst*, *Defb*, *Wfdc*, *Spint*, *Spink*, *Serpin*) across the fifteen principal cell subclusters from **Figure 3B**. B) Three examples of relevant multigene clusters exhibiting region-specific expression of individual cluster members.

**Figure 3-figure supplement 2.**
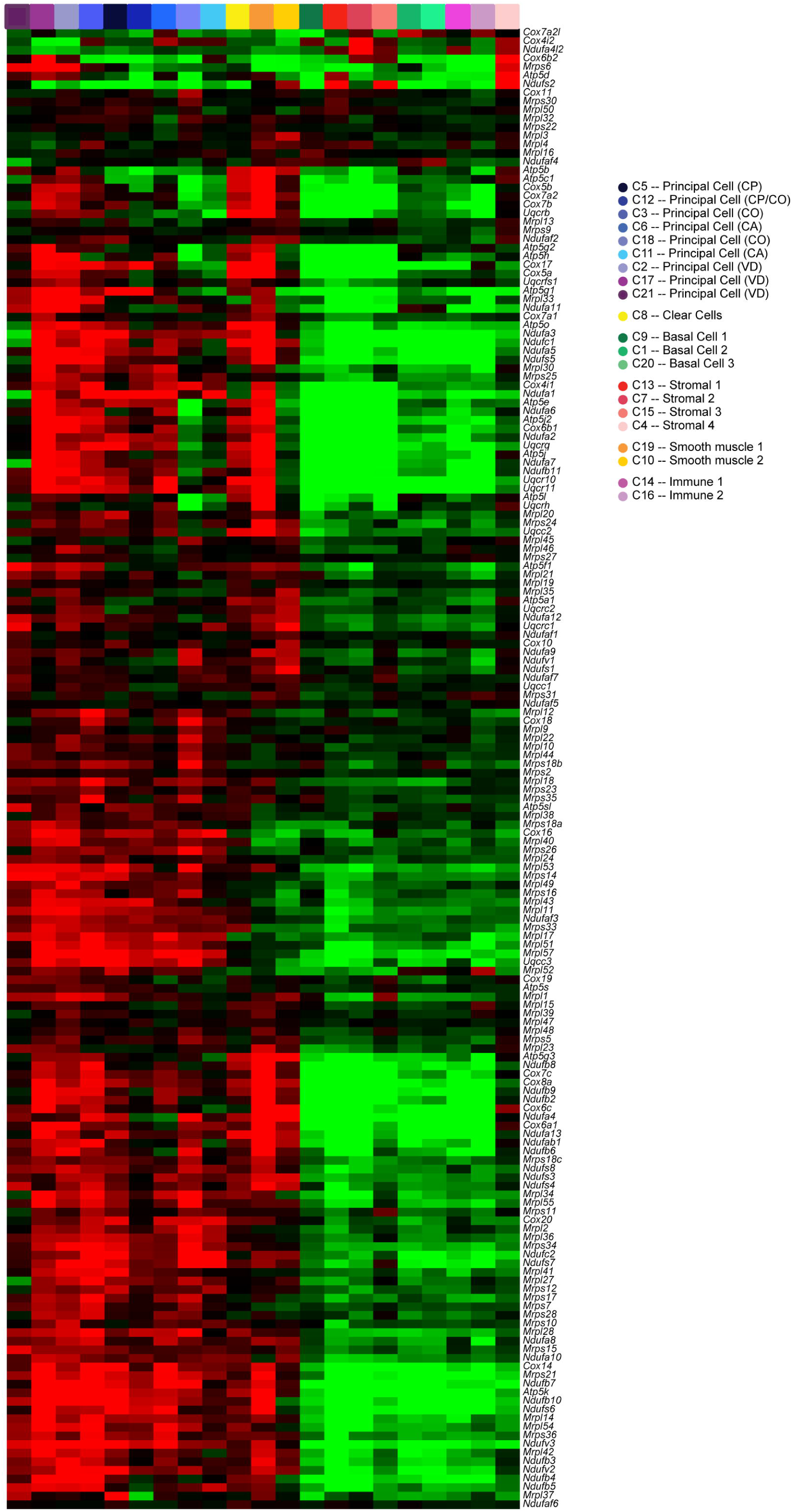
High levels of oxidative energy production in principal, clear, and muscle cells. Heatmap shows expression of various nuclear-encoded mitochondrial genes (*Mrpl*, *Mrps*, *Ndufa*, *Cox*, *Uqcr* gene families) across the entire 21 cluster dataset from **Figure 2A**. Clusters expressing high levels of nuclear-encoded mitochondrial genes included all principal cell clusters, both muscle clusters, and the clear cells (leftmost 12 columns), while low expression of these genes was observed in basal, immune, and stromal cell types (rightmost 9 columns).

**Figure 3-figure supplement 3.**
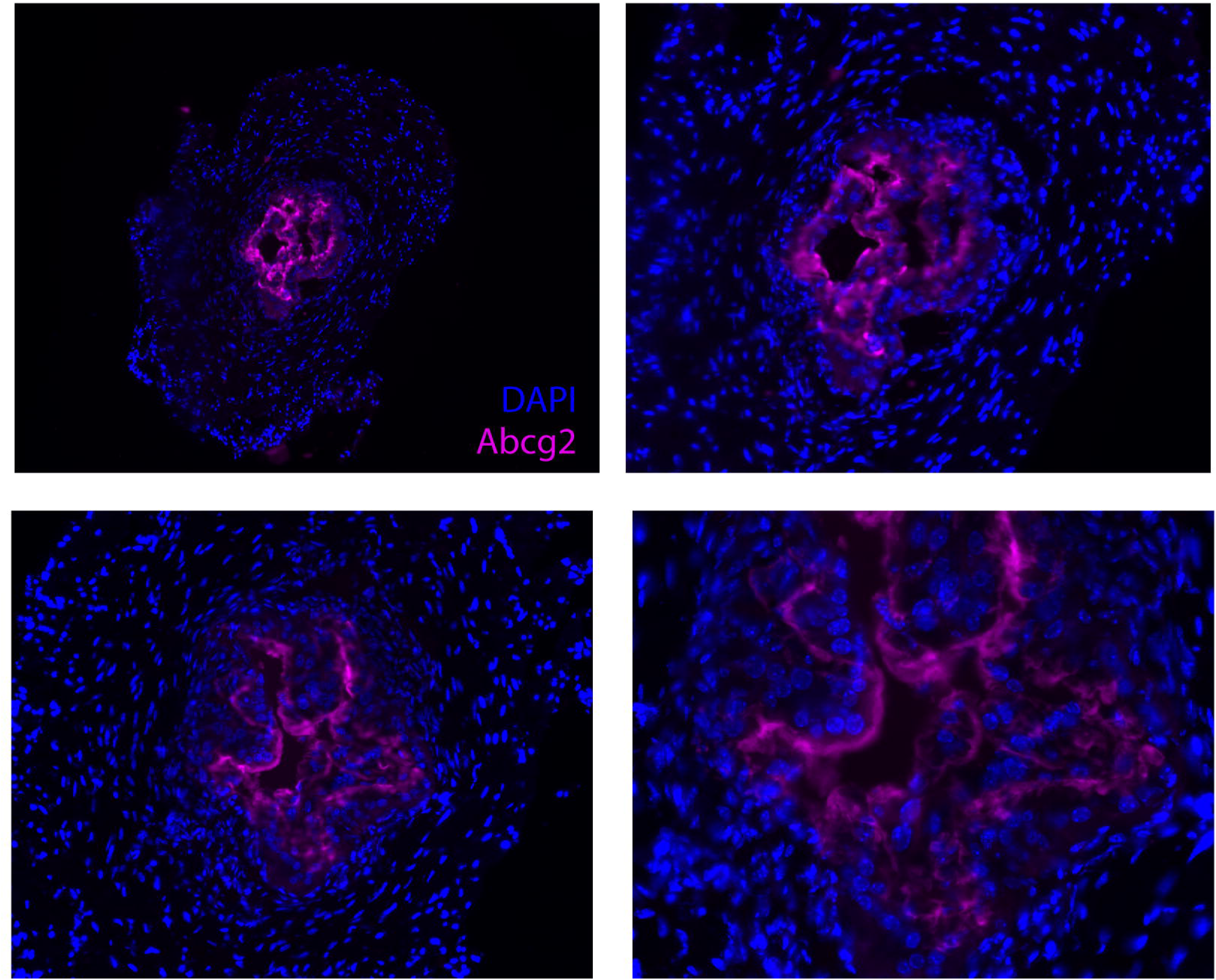
Apical localization of Abcg2 in the vas deferens. Immunostaining for Abcg2, counterstained with DAPI, in the murine vas deferens.

**Figure 5-figure supplement 1.**
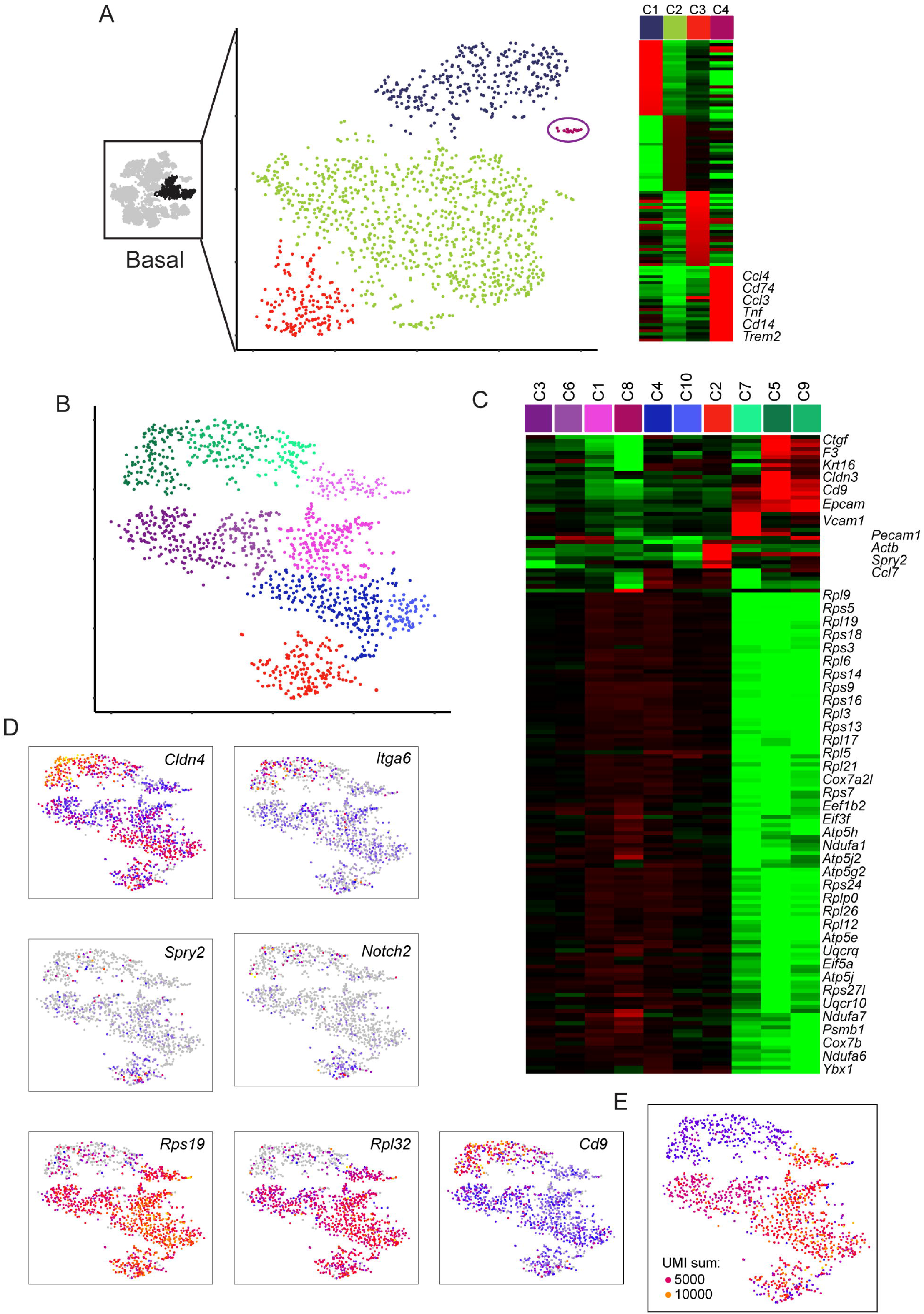
Subclustering of basal cells. A) Initial subclustering of basal cells. Left panel highlights the three clusters from the full dataset used for reclustering, which is shown in the next panel. However, this subclustering revealed a small and highly distinct population (∼15 cells) of macrophages (markers shown in right panel). These cells were removed for subsequent basal cell reclustering. B) Reclustering of basal cells with macrophages removed, with 10 clusters annotated by color. C) Expression of marker genes (minimum fold-enrichment > 4-fold in at least one cluster) across the ten new basal subclusters, grouped by hierarchical clustering. Notable here are two major populations, distinguished by expression of ribosomal protein genes. A third minor cluster (C2) was marked by elevated expression of several genes including *Notch2* and *Spry2*. D) Expression of individual genes across the basal cell subclusters, as in **Figure 2C**. E) UMI counts for the basal cell subclustering, revealing a strong correspondence between the RPG high/low divisions and cells with high/low UMI counts.

**Figure 5-figure supplement 2.**
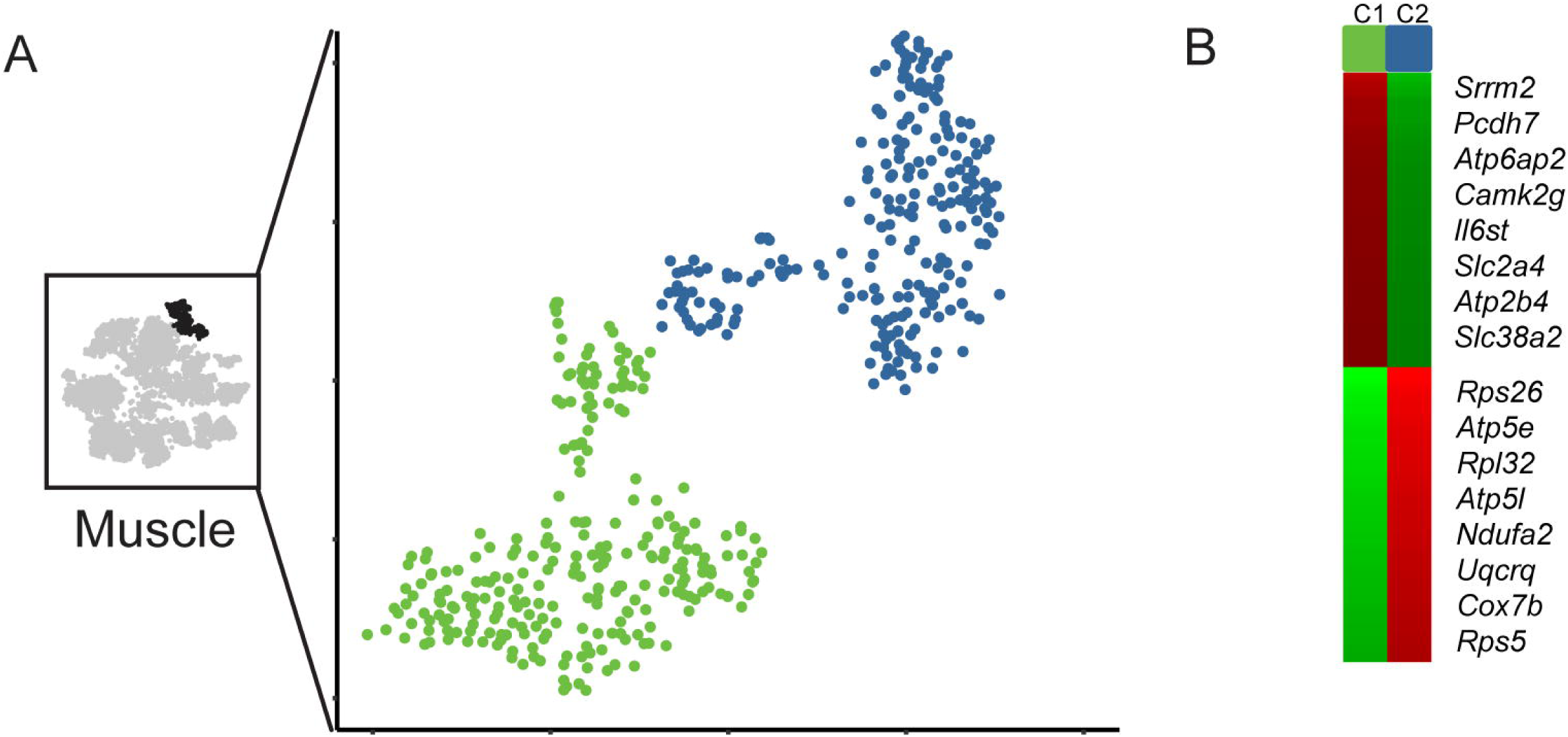
Subclustering of muscle cells. A) Subclustering of muscle cells. Panels arranged similarly to **Figure5-figure supplement 1A**. B) Markers distinguishing the two muscle cell subtypes.

## REFERENCES

Balhorn, R. (1982). A model for the structure of chromatin in mammalian sperm. The Journal of cell biology 93, 298–305.

Bedford, J.M. (1967). Effects of duct ligation on the fertilizing ability of spermatozoa from different regions of the rabbit epididymis. J Exp Zool 166, 271–281.

Bjorkgren, I., Saastamoinen, L., Krutskikh, A., Huhtaniemi, I., Poutanen, M., and Sipila, P. (2012). Dicer1 ablation in the mouse epididymis causes dedifferentiation of the epithelium and imbalance in sex steroid signaling. PloS one 7, e38457.

Boyle, C.N., and Le Foll, C. (2019). Amylin and Leptin interaction: Role During Pregnancy, Lactation and Neonatal Development. Neuroscience.

Breton, S., Hammar, K., Smith, P.J., and Brown, D. (1998). Proton secretion in the male reproductive tract: involvement of Cl--independent HCO-3 transport. Am J Physiol 275, C1134–1142.

Breton, S., Ruan, Y.C., Park, Y.J., and Kim, B. (2016). Regulation of epithelial function, differentiation, and remodeling in the epididymis. Asian journal of andrology 18, 3–9.

Breton, S., Smith, P.J., Lui, B., and Brown, D. (1996). Acidification of the male reproductive tract by a proton pumping (H+)-ATPase. Nature medicine 2, 470–472.

Breton, S., Tyszkowski, R., Sabolic, I., and Brown, D. (1999). Postnatal development of H+ ATPase (proton-pump)-rich cells in rat epididymis. Histochem Cell Biol 111, 97–105.

Brown, D., Lui, B., Gluck, S., and Sabolic, I. (1992). A plasma membrane proton ATPase in specialized cells of rat epididymis. Am J Physiol 263, C913–916.

Caballero, J., Frenette, G., D’Amours, O., Dufour, M., Oko, R., and Sullivan, R. (2012). ATP-binding cassette transporter G2 activity in the bovine spermatozoa is modulated along the epididymal duct and at ejaculation. Biology of reproduction 86, 181.

Caballero, J.N., Frenette, G., Belleannee, C., and Sullivan, R. (2013). CD9-positive microvesicles mediate the transfer of molecules to Bovine Spermatozoa during epididymal maturation. PloS one 8, e65364.

Calvin, H.I., and Bedford, J.M. (1971). Formation of disulphide bonds in the nucleus and accessory structures of mammalian spermatozoa during maturation in the epididymis. J Reprod Fertil Suppl 13, Suppl 13:65–75.

Cao, J., Packer, J.S., Ramani, V., Cusanovich, D.A., Huynh, C., Daza, R., Qiu, X., Lee, C., Furlan, S.N., Steemers, F.J., et al. (2017). Comprehensive single-cell transcriptional profiling of a multicellular organism. Science (New York, NY 357, 661–667.

Carr, D.W., and Acott, T.S. (1984). Inhibition of bovine spermatozoa by caudal epididymal fluid: I. Studies of a sperm motility quiescence factor. Biology of reproduction 30, 913–925.

Chen, Q., Yan, M., Cao, Z., Li, X., Zhang, Y., Shi, J., Feng, G.H., Peng, H., Zhang, X., Zhang, Y., et al. (2016). Sperm tsRNAs contribute to intergenerational inheritance of an acquired metabolic disorder. Science (New York, NY 351, 397–400.

Clauss, A., Persson, M., Lilja, H., and Lundwall, A. (2011). Three genes expressing Kunitz domains in the epididymis are related to genes of WFDC-type protease inhibitors and semen coagulum proteins in spite of lacking similarity between their protein products. BMC Biochem 12, 55.

Cohen, M., Giladi, A., Gorki, A.D., Solodkin, D.G., Zada, M., Hladik, A., Miklosi, A., Salame, T.M., Halpern, K.B., David, E., et al. (2018). Lung Single-Cell Signaling Interaction Map Reveals Basophil Role in Macrophage Imprinting. Cell 175, 1031–1044 e1018.

Conine, C.C., Sun, F., Song, L., Rivera-Perez, J.A., and Rando, O.J. (2018). Small RNAs Gained during Epididymal Transit of Sperm Are Essential for Embryonic Development in Mice. Developmental cell 46, 470–480 e473.

Cooper, T.G. (2015). Epididymal research: more warp than weft? Asian journal of andrology 17, 699–703.

Da Silva, N., and Barton, C.R. (2016). Macrophages and dendritic cells in the post-testicular environment. Cell Tissue Res 363, 97–104.

Derr, A., Yang, C., Zilionis, R., Sergushichev, A., Blodgett, D.M., Redick, S., Bortell, R., Luban, J., Harlan, D.M., Kadener, S., et al. (2016). End Sequence Analysis Toolkit (ESAT) expands the extractable information from single-cell RNA-seq data. Genome research 26, 1397–1410.

Domeniconi, R.F., Souza, A.C., Xu, B., Washington, A.M., and Hinton, B.T. (2016). Is the Epididymis a Series of Organs Placed Side By Side? Biology of reproduction 95, 10.

Farrell, J.A., Wang, Y., Riesenfeld, S.J., Shekhar, K., Regev, A., and Schier, A.F. (2018). Single-cell reconstruction of developmental trajectories during zebrafish embryogenesis. Science (New York, NY 360.

Gervasi, M.G., and Visconti, P.E. (2017). Molecular changes and signaling events occurring in spermatozoa during epididymal maturation. Andrology 5, 204–218.

Golonka, R., Yeoh, B.S., and Vijay-Kumar, M. (2019). The Iron Tug-of-War between Bacterial Siderophores and Innate Immunity. J Innate Immun, 1–14.

Gregory, M., and Cyr, D.G. (2014). The blood-epididymis barrier and inflammation. Spermatogenesis 4, e979619.

Guyonnet, B., Marot, G., Dacheux, J.L., Mercat, M.J., Schwob, S., Jaffrezic, F., and Gatti, J.L. (2009). The adult boar testicular and epididymal transcriptomes. BMC Genomics 10, 369.

Hall, S.H., Yenugu, S., Radhakrishnan, Y., Avellar, M.C., Petrusz, P., and French, F.S. (2007). Characterization and functions of beta defensins in the epididymis. Asian journal of andrology 9, 453–462.

Han, X., Wang, R., Zhou, Y., Fei, L., Sun, H., Lai, S., Saadatpour, A., Zhou, Z., Chen, H., Ye, F., et al. (2018). Mapping the Mouse Cell Atlas by Microwell-Seq. Cell 172, 1091–1107 e1017.

Hsia, N., and Cornwall, G.A. (2004). DNA microarray analysis of region-specific gene expression in the mouse epididymis. Biology of reproduction 70, 448–457.

Jagoe, W.N., Howe, K., O’Brien, S.C., and Carroll, J. (2013). Identification of a role for a mouse sperm surface aldo-keto reductase (AKR1B7) and its human analogue in the detoxification of the reactive aldehyde, acrolein. Andrologia 45, 326–331.

Jelinsky, S.A., Turner, T.T., Bang, H.J., Finger, J.N., Solarz, M.K., Wilson, E., Brown, E.L., Kopf, G.S., and Johnston, D.S. (2007). The rat epididymal transcriptome: comparison of segmental gene expression in the rat and mouse epididymides. Biology of reproduction 76, 561–570.

Jervis, K.M., and Robaire, B. (2001). Dynamic changes in gene expression along the rat epididymis. Biology of reproduction 65, 696–703.

Johnston, D.S., Jelinsky, S.A., Bang, H.J., DiCandeloro, P., Wilson, E., Kopf, G.S., and Turner, T.T. (2005). The mouse epididymal transcriptome: transcriptional profiling of segmental gene expression in the epididymis. Biology of reproduction 73, 404–413.

Kabututu, Z., Manin, M., Pointud, J.C., Maruyama, T., Nagata, N., Lambert, S., Lefrancois-Martinez, A.M., Martinez, A., and Urade, Y. (2009). Prostaglandin F2alpha synthase activities of aldo-keto reductase 1B1, 1B3 and 1B7. J Biochem 145, 161–168.

Kim, D., Paggi, J.M., Park, C., Bennett, C., and Salzberg, S.L. (2019). Graph-based genome alignment and genotyping with HISAT2 and HISAT-genotype. Nature biotechnology 37, 907–915.

Krapf, D., Ruan, Y.C., Wertheimer, E.V., Battistone, M.A., Pawlak, J.B., Sanjay, A., Pilder, S.H., Cuasnicu, P., Breton, S., and Visconti, P.E. (2012). cSrc is necessary for epididymal development and is incorporated into sperm during epididymal transit. Developmental biology 369, 43–53.

Lareyre, J.J., Zheng, W.L., Zhao, G.Q., Kasper, S., Newcomer, M.E., Matusik, R.J., Ong, D.E., and Orgebin-Crist, M.C. (1998). Molecular cloning and hormonal regulation of a murine epididymal retinoic acid-binding protein messenger ribonucleic acid. Endocrinology 139, 2971–2981.

Lun, A.T., Bach, K., and Marioni, J.C. (2016). Pooling across cells to normalize single-cell RNA sequencing data with many zero counts. Genome biology 17, 75.

Ma, L., Yu, H., Ni, Z., Hu, S., Ma, W., Chu, C., Liu, Q., and Zhang, Y. (2013). Spink13, an epididymis-specific gene of the Kazal-type serine protease inhibitor (SPINK) family, is essential for the acrosomal integrity and male fertility. The Journal of biological chemistry 288, 10154–10165.

Mandon, M., Hermo, L., and Cyr, D.G. (2015). Isolated Rat Epididymal Basal Cells Share Common Properties with Adult Stem Cells. Biology of reproduction 93, 115.

Martin-DeLeon, P.A. (2015). Epididymosomes: transfer of fertility-modulating proteins to the sperm surface. Asian journal of andrology 17, 720–725.

Martinez, G., Choudury, S.G., and Slotkin, R.K. (2017). tRNA-derived small RNAs target transposable element transcripts. Nucleic acids research 45, 5142–5152.

McCarthy, D.J., Chen, Y., and Smyth, G.K. (2012). Differential expression analysis of multifactor RNA-Seq experiments with respect to biological variation. Nucleic acids research 40, 4288–4297.

Oliveira, R.L., Parent, A., Cyr, D.G., Gregory, M., Mandato, C.A., Smith, C.E., and Hermo, L. (2016). Implications of caveolae in testicular and epididymal myoid cells to sperm motility. Molecular reproduction and development 83, 526–540.

Ong, D.E., Newcomer, M.E., Lareyre, J.J., and Orgebin-Crist, M.C. (2000). Epididymal retinoic acid-binding protein. Biochimica et biophysica acta 1482, 209–217.

Orgebin-Crist, M.C. (1967). Sperm maturation in rabbit epididymis. Nature 216, 816–818.

Ramilowski, J.A., Goldberg, T., Harshbarger, J., Kloppmann, E., Lizio, M., Satagopam, V.P., Itoh, M., Kawaji, H., Carninci, P., Rost, B., et al. (2015). A draft network of ligand-receptor-mediated multicellular signalling in human. Nature communications 6, 7866.

Ramskold, D., Luo, S., Wang, Y.C., Li, R., Deng, Q., Faridani, O.R., Daniels, G.A., Khrebtukova, I., Loring, J.F., Laurent, L.C., et al. (2012). Full-length mRNA-Seq from single-cell levels of RNA and individual circulating tumor cells. Nature biotechnology 30, 777–782.

Reilly, J.N., McLaughlin, E.A., Stanger, S.J., Anderson, A.L., Hutcheon, K., Church, K., Mihalas, B.P., Tyagi, S., Holt, J.E., Eamens, A.L., et al. (2016). Characterisation of mouse epididymosomes reveals a complex profile of microRNAs and a potential mechanism for modification of the sperm epigenome. Sci Rep 6, 31794.

Rodriguez, A., and Laio, A. (2014). Machine learning. Clustering by fast search and find of density peaks. Science (New York, NY 344, 1492–1496.

Rompala, G.R., Mounier, A., Wolfe, C.M., Lin, Q., Lefterov, I., and Homanics, G.E. (2018). Heavy Chronic Intermittent Ethanol Exposure Alters Small Noncoding RNAs in Mouse Sperm and Epididymosomes. Front Genet 9, 32.

Saez, F., Ouvrier, A., and Drevet, J.R. (2011). Epididymis cholesterol homeostasis and sperm fertilizing ability. Asian journal of andrology 13, 11–17.

Schorn, A.J., Gutbrod, M.J., LeBlanc, C., and Martienssen, R. (2017). LTR-Retrotransposon Control by tRNA-Derived Small RNAs. Cell 170, 61–71 e11.

Serre, V., and Robaire, B. (1999). Distribution of immune cells in the epididymis of the aging Brown Norway rat is segment-specific and related to the luminal content. Biology of reproduction 61, 705–714.

Sharma, U., Conine, C.C., Shea, J.M., Boskovic, A., Derr, A.G., Bing, X.Y., Belleannee, C., Kucukural, A., Serra, R.W., Sun, F., et al. (2016). Biogenesis and function of tRNA fragments during sperm maturation and fertilization in mammals. Science (New York, NY 351, 391–396.

Sharma, U., Sun, F., Conine, C.C., Reichholf, B., Kukreja, S., Herzog, V.A., Ameres, S.L., and Rando, O.J. (2018). Small RNAs Are Trafficked from the Epididymis to Developing Mammalian Sperm. Developmental cell 46, 481–494 e486.

Shum, W.W., Da Silva, N., Belleannee, C., McKee, M., Brown, D., and Breton, S. (2011). Regulation of V-ATPase recycling via a RhoA- and ROCKII-dependent pathway in epididymal clear cells. Am J Physiol Cell Physiol 301, C31–43.

Shum, W.W., Da Silva, N., McKee, M., Smith, P.J., Brown, D., and Breton, S. (2008). Transepithelial projections from basal cells are luminal sensors in pseudostratified epithelia. Cell 135, 1108–1117.

Sullivan, R., Frenette, G., and Girouard, J. (2007). Epididymosomes are involved in the acquisition of new sperm proteins during epididymal transit. Asian journal of andrology 9, 483–491.

Sullivan, R., and Saez, F. (2013). Epididymosomes, prostasomes, and liposomes: their roles in mammalian male reproductive physiology. Reproduction 146, R21–35.

Sun, E.L., and Flickinger, C.J. (1979). Development of cell types and of regional differences in the postnatal rat epididymis. Am J Anat 154, 27–55.

Sun, E.L., and Flickinger, C.J. (1980). Morphological characteristics of cells with apical nuclei in the initial segment of the adult rat epididymis. Anat Rec 196, 285–293.

Tecle, E., and Gagneux, P. (2015). Sugar-coated sperm: Unraveling the functions of the mammalian sperm glycocalyx. Molecular reproduction and development 82, 635–650.

Thimon, V., Koukoui, O., Calvo, E., and Sullivan, R. (2007). Region-specific gene expression profiling along the human epididymis. Mol Hum Reprod 13, 691–704.

Tulsiani, D.R. (2006). Glycan-modifying enzymes in luminal fluid of the mammalian epididymis: an overview of their potential role in sperm maturation. Mol Cell Endocrinol 250, 58–65.

Turner, T.T., Bomgardner, D., Jacobs, J.P., and Nguyen, Q.A. (2003). Association of segmentation of the epididymal interstitium with segmented tubule function in rats and mice. Reproduction 125, 871–878.

van der Maaten, L.J.P., and Hinton, G.E. (2008). Visualizing High-Dimensional Data Using t-SNE. Journal of Machine Learning Research 9, 2579–2605.

Volat, F.E., Pointud, J.C., Pastel, E., Morio, B., Sion, B., Hamard, G., Guichardant, M., Colas, R., Lefrancois-Martinez, A.M., and Martinez, A. (2012). Depressed levels of prostaglandin F2alpha in mice lacking Akr1b7 increase basal adiposity and predispose to diet-induced obesity. Diabetes 61, 2796–2806.

Wang, H., Wu, X., Hudkins, K., Mikheev, A., Zhang, H., Gupta, A., Unadkat, J.D., and Mao, Q. (2006). Expression of the breast cancer resistance protein (Bcrp1/Abcg2) in tissues from pregnant mice: effects of pregnancy and correlations with nuclear receptors. American journal of physiology Endocrinology and metabolism 291, E1295–1304.

Young, W.C. (1931). A study of the function of the epididymis. III. Functional changes undergone by spermatozoa during their passage through the epididymis and vas deferens in the guinea pig. J Exp Biol 8, 151–162.

